# p70 ribosomal protein S6 Kinase (p70S6K) as a potential peripheral biomarker for autistic symptoms in 16p11.2 deletion and duplication syndromes

**DOI:** 10.64898/2026.02.23.706592

**Authors:** Ilaria Morella, Jessica Hall, Charlotte Butter, Caitlin Goldie, Georgia Evans, Nabila Ali, Natalia Burlinson-Diaz, Emma Burkitt-Wright, Isabella Caraiscos, Lorenzo Morè, Jonathan Green, Marianne B.M. van den Bree, Riccardo Brambilla

## Abstract

Neurodevelopmental disorders (NDDs) encompass cognitive, motor, and psychiatric manifestations that require extensive assessments spanning different clinical domains. Individuals with syndromes caused by micro-deletions and -duplications in the proximal region of 16p11.2 have increased likelihood of neurodevelopmental and mental health symptoms, as well as cardiometabolic disorder. However, considerable phenotypic variability complicates prediction of risk, disease trajectory and treatment response. The identification of biomarkers in easily accessible tissues, such as Peripheral Blood Mononuclear Cells (PBMCs), could help diagnosis and prognosis, yet remains an unmet clinical need. Converging evidence implicates dysregulation of ubiquitous signalling pathways, including ERK and mTOR cascades, in altered protein synthesis across NDDs of both idiopathic and genetic origin. Here, using a hypothesis-driven approach, we examined candidate peripheral biomarkers in individuals with 16p11.2 proximal deletion and duplication syndromes. We established increased psychopathology and cognitive deficits in individuals with 16p11.2 deletion and duplication syndromes, compared to sibling controls. Furthermore, we identified a significant reduction of p70 S6 kinase (p70S6K) levels in PBMCs across both genotypes compared to controls. Notably, the magnitude of p70S6K reduction correlated with the severity of autistic symptoms independent of CNV genotype. In contrast, MAPK3/ERK1 protein levels mirrored the gene dosage, showing increased expression in individuals with the duplication and reduced levels in those with the deletion. These findings suggest that peripheral alterations in p70S6K may relate to variation in autism-related traits within individuals with 16p11.2 deletion and duplication syndromes, indicating that it may represent a potential biomarker of autism-associated symptom severity in this genetically at-risk group.

## Introduction

Copy number variants (CNVs) at the 16p11.2 locus, specifically the recurrent deletion and duplication of genetic material spanning breakpoints 4 and 5 (BP4-BP5) (593 kb; chr16; 29.6–30.2 mb-HG19), are prevalent in approximately 1/2000 and 1/1100 births respectively, in the general population . Both 16p11.2 deletion and duplication syndromes contribute to highly complex and heterogeneous phenotypic presentations, which are characterised by variable expressivity. Studies have reported increased risk of emotional and behavioural difficulties (2) and neuropsychiatric and neurodevelopmental symptoms, including intellectual disability (ID) (3, 4), development coordination difficulties (5), autism spectrum disorder (ASD) (3, 6, 7), attention deficit hyperactivity disorder (ADHD) (3, 8), mood and anxiety disorders (8), and epilepsy (9). Individuals with 16p11.2 duplication syndrome tend to exhibit more severe symptoms than those with 16p11.2 deletion syndrome, including increased risk of any psychiatric condition, psychotic symptoms and worse cognitive performance (3, 8, 10, 11). Both conditions are associated with metabolic and cardiovascular symptoms (12).

The 16p11.2 proximal locus contains 27 protein-coding genes, including signalling modulators, transcription factors and chromatin remodellers (11). Evidence in experimental models suggests that multiple genes within the locus may interact via interlinked molecular pathways contributing to the phenotypic variability, but the exact neurobiological mechanisms underlying their clinical phenotypes remain unknown (11, 13).

The Mitogen Activated Protein Kinase 3 (MAPK3), encoding the Extracellular Signal-Regulated Kinase 1 (ERK1), is located within the 16p11.2 locus. ERK1 is closely related with ERK2, encoded by MAPK1, a gene located on chromosome 22. ERK1 and ERK2 form a core component of the Ras-ERK signalling pathway which plays key roles in cell proliferation, differentiation, survival, synaptogenesis, and behavioural plasticity (14, 15). The Ras-ERK pathway is initiated upon a broad range of extracellular stimuli, including neurotransmitters, growth factors, calcium and morphogens. Activated Ras triggers the phosphorylation of Raf-MEK-ERK cascade, ultimately resulting in the activation of transcription factors and cytoplasmic targets (16). Due to its essential role during neurodevelopment, disruptions in the Ras-ERK pathway contribute to the neuropathology of several neurodevelopmental disorders (NDDs), including ASD, ADHD, Fragile X syndrome, schizophrenia (17) and “RASopathies” (18). In a 16p11.2 deletion mouse model, reduced ERK1 expression causes ERK signalling hyperactivation, associated with defects in cortical neurogenesis, ultimately leading to aberrant cortical circuitry and behavioural dysfunctions, including hyperactivity, impaired hippocampal-based memory function, anxiety, deficits in olfaction and impaired maternal behaviour (19, 20). Interestingly, pre-natal administration of cell permeable peptides inhibiting ERK pathway rescued deficits in neurogenesis and completely restored hippocampal-based memory, anxiety, olfaction, maternal behaviour and hyperactivity in 16p11.2 deletion mice, while post-natal treatment partially ameliorates these deficits (19).

The ERK1/2 pathway converges onto the phosphatidylinositol 3-kinase-AKT-mechanistic target of the rapamycin (PI3K-AKT-mTOR) signalling pathway (21), which has prominent roles in local protein synthesis at the synapses (16) and in brain development (21). The mTOR kinase exists in two functional complexes, mTOR Complex 1 (mTORC1) and mTOR Complex 2 (mTORC2). The PI3K-AKT-mTOR is activated by tyrosine kinase receptors and metabotropic receptors, leading to the activation of AKT which disinhibits mTOR by blocking the Tuberous Sclerosis Complex 1 and 2 (TSC1 and 2). mTORC1 promotes protein synthesis by phosphorylating p70 ribosomal protein S6 kinase (p70S6K), which acts on downstream substrates involved in translation initiation and elongation. In addition, mTORC1-mediated phosphorylation of the eukaryotic initiation factor 4E (eIF4E)-binding protein (4E-BP) allows the assembly of eIF4E mRNA cap-binding protein into the cap-binding complex to initiate translation (16). Mutations of components of the PI3K-AKT-mTOR pathway (including TSC1, eIF4E and 4EBP2) result in translational dysregulation leading to synaptic plasticity defects and have been associated with syndromic forms of ASD and ID (18). However, the role of the PI3K-AKT-mTOR pathway in 16p11.2 CNVs has not been established yet.

This work aims to fill this gap in the literature. This study aimed to: 1) establish whether 16p11.2 CNV status confers increased risk for adverse neurodevelopmental, mental health, and cognitive outcomes in a clinical cohort of children and young people (CYP) and, secondarily, to examine whether profiles differ between 16p11.2 deletion and duplication, 2) determine the levels of multiple signalling proteins along the ERK and mTOR pathway in CYP with 16p11.2 deletion and duplication compared to sibling controls, and 3) investigate whether alterations in peripheral signalling proteins are associated with the severity of neurodevelopmental, mental health and cognitive outcomes. Finally, we conducted exploratory mediation analyses, where supported by findings from aims 1–3, to examine potential mechanisms. This approach may establish biological markers associated with neurodevelopmental, mental health and cognitive outcomes in individuals with 16p11.2 deletion and duplication syndromes and may inform future pharmacological studies, with potential implications beyond the 16p11.2 deletion and duplication syndromes.

## Methods

### Study participants

CYP with proximal 16p11.2 deletion syndrome (n=45) or duplication syndrome (n=29) were recruited by teams at Cardiff University and the University of Manchester, via Medical Genetics clinics in the UK, charities (Unique, MaxAppeal), social media and word of mouth. Where possible, a sibling, closest in age to the CYP with 16p11.2 deletion/duplication syndrome, and who did not have a known pathogenic CNV, was also recruited. They comprised our control group (n=30). Participant demographics are presented in **Table 1**.

**Table 1.** Demographic characteristics are presented for CYP with 16p11.2 deletion and duplication, and control participants. Values are reported as *n* (%) for categorical variables and mean (**±**SD) for age. Maternal ethnicity was used as the indicator of ethnic ancestry. Maternal education and household income were based on parent-reported measures. Control participants were recruited from the same households as CYP with 16p11.2 deletion and duplication and therefore shared family environmental characteristics. Sample sizes (*n*) for each group are reported in the table.

|  | Deletion | Duplication | Controls | Statistics (X<sup>2</sup> and ANOVA) | Pairwise Comparisons (BH-FDR corrected) |  |  |
| --- | --- | --- | --- | --- | --- | --- | --- |
|  | n = 45 | n = 29 | n= 30 |  | Deletion-control | Duplication-control | Deletion-Duplication |
| Mean Age (±SD) | 10.3 (±2.4) | 11.2 (±3.0) | 12.4 (±2.8) | F=6.008, df=2, p=0.005 | p=0.003 | p=0.148 | p=0.148 |
| Male | 23 (51.1%) | 19 (65.5%) | 8 (26.7%) | X<sup>2</sup>=9.208, df=2, p=0.010 | p=0.052 | p=0.008 | p=0.222 |
|  | Deletion |  |  | Duplication |  |  |  |
|  | n=45 |  |  | n=29 |  |  |  |
| Ethnic Ancestry |  |  |  |  |  |  |  |
| White British | 42 (93.3%) |  |  | 28 (96.6%) |  |  |  |
| Other | 3 (6.7%) |  |  | 1 (3.4%) |  |  |  |
| Mothers Education |  |  |  |  |  |  |  |
| Low (<9 years) | 3 (6.7%) |  |  | 3 (10.3%) |  |  |  |
| Average (A levels or equivalent) | 27 (60%) |  |  | 20 (69.0%) |  |  |  |
| High (higher education) | 14 (31.1%) |  |  | 6 (20.7%) |  |  |  |
| Missing | 1 (2.2%) |  |  | NA |  |  |  |
| Approx household income |  |  |  |  |  |  |  |
| Low (<£20,000) | 11 (24.5%) |  |  | 9 (31.1%) |  |  |  |
| Average (£20-39,999) | 18 (40%) |  |  | 11 (37.9%) |  |  |  |
| High (£40,000+) | 15 (33.3%) |  |  | 8 (27.6%) |  |  |  |
| Refused | 1 (2.2%) |  |  | 1 (3.4%) |  |  |  |

Saliva samples were collected from all participants and genotyped at Cardiff University using microarray techniques (Illumina Global Screening Array (GSAv3.0)). Methodological details are provided in **Supplementary materials and methods**.

Written consent was obtained from the primary caregiver, and consent or assent by the CYP, as applicable. All procedures contributing to this work comply with the ethical standards of the relevant national and institutional committees on human experimentation and with the Helsinki Declaration of 1975, as revised in 2013. All procedures involving human participants were approved by the lead university’s research ethics committee - Cardiff School of Medicine, the National Health Service (NHS) East Midlands - Leicester Central Research Ethics Committee (19/EM/0287) and by all NHS ethics and research and development committees of participating UK Clinical Genetics Clinics.

### Assessments

Data were collected by trained researchers either in person at participants’ homes, or onsite at the Royal Manchester Children’s Hospital (RMCH), or the Cardiff University Division of Psychological Medicine and Clinical Neurosciences (DPMCN). The study ran from September 2020 to September 2024, during which the COVID-19 pandemic took place. Due to restrictions during this period, phenotypic assessments were also conducted remotely where necessary. Saliva samples were obtained in person or by post (when assessments were conducted remotely), while blood samples were collected at RMCH under strict clinical protocols, with play therapist support when needed. Blood samples were only acquired from a subset of the sample, who were able to provide one (n = 18 controls, 23 deletion, 15 duplication).

Parents/caregivers completed the Child and Adolescent Psychiatric Assessment (CAPA) (19), a semi-structured interview yielding symptom counts for all psychiatric conditions, except autism. Focus areas included ADHD, ODD, OCD, mood disorders (depression, mania, hypomania), and anxiety disorders (e.g., GAD, social/specific phobia, separation anxiety, selective mutism, panic disorder).

Autistic traits were screened using the Social Communication Questionnaire (SCQ) (20). CYP scoring >15 on the SCQ, indicative of potential ASD (22) were invited for further assessment via the Autism Diagnostic Interview-Revised (ADI-R (23)), an in-depth, semi-structured caregiver interview that evaluates autistic symptoms across social interaction, communication, and restricted/repetitive behaviour domains. It generates algorithm scores aligned with DSM-5 diagnostic criteria for autism and a total count of autistic symptoms.

The Developmental Coordination Disorder Questionnaire (DCDQ) (22) was used to screen for Developmental Coordination Disorder (DCD). The DCDQ is a caregiver-report questionnaire consisting of 15 items rated on a 5-point likert scale that assess a child’s everyday motor performances compared to peers, covering: control during movement, fine motor/handwriting, and general coordination. As DCDQ alone cannot establish a formal research diagnosis, DCDQ was used to identify “indicative DCD”. In the DCDQ, lower scores indicate greater motor coordination difficulties, and the age-specific cut-offs identify children whose scores fall below a threshold consistent with “indicative DCD.” Specifically, children aged 5–7 years 11 months with scores of 15–46, those aged 8–9 years 11 months with scores of 15–55, and those aged 10 years and above with scores of 15–57 meet criteria for “indicative DCD”. To allow direct comparison with other diagnostic measures where *higher* scores reflect greater impairment, simplify interpretation and maintain conceptual alignment across diagnostic categories, DCDQ total scores were inverted.

Intelligence quotient (IQ) was assessed using the Wechsler Abbreviated Scale of Intelligence (WASI), comprising verbal (VIQ) and performance (PIQ) subtests, with composite and full-scale IQ (FSIQ) scores calculated. During the COVID-19 pandemic, the block design PIQ subtest was unavailable for online administration, due to inability to carry out manual manipulation of the blocks. Thus, only VIQ and FSIQ scores were available. FSIQ scores from both two- and four-subtest versions were treated equivalently. An intellectual disability variable (ID) was derived, where an FSIQ ≤70 was deemed to be representative of ID.

An “any psychiatric disorder” variable was derived, whereby any psychiatric disorder was inclusive of ADHD, anxiety, ODD, OCD, mood disorder, or ASD (SCQ ≥15).

### Isolation of Peripheral Blood Mononuclear Cells (PBMCs) and protein extraction

Following phenotypic assessments, blood samples (8-9 ml per participant, with up to 20ml when feasible) were collected by venepuncture using EDTA-coated tubes. Immediately after blood collection, PBMCs were isolated by density gradient using Accuspin Histopaque-1077 system (Sigma Aldrich), following the manufacturer’s instructions. Cell pellets were resuspended in 50 μl of ice-cold lysis buffer (20 mM Tris HCl pH 7.5, 150 mM NaCl, 1 mM EDTA, 1% Triton X-100, 1X cOmplete^®^ EDTA-free protease inhibitor cocktail (Roche), 1 mM sodium orthovanadate, 1 mM glycerophosphate, 1 mM sodium fluoride). Protein concentration was determined using the Biorad DC Protein assay.

### Automated Western blotting assay

Automated Western blotting was performed using Jess Simple Western system (Protein Simple), following the manufacturer’s method for the 12-230-kDa Jess separation module.

Protein lysates were mixed with 0.1X Sample Buffer (Protein Simple) and Fluorescent 5X Master Mix (Protein Simple) to reach a concentration of 1 μg/μl (ERK1/2, phospho-ERK1/2, phospho-eIF4E and TSC1) or 2 μg/μl (eIF4E, AKT, phospho-AKT, p70S6K and phospho-p70S6K), and denatured at 95°C for 5 min. A proprietary normalization reagent (Protein Simple) was used to normalize over total protein levels in each sample. Following electrophoresis, primary antibodies were applied for 1 h after a blocking step (details provided in **Supplementary Table 1**). HRP-conjugated anti-mouse or anti-rabbit antibodies (Protein Simple) were applied for 30 min, followed by peroxide/luminol-S (Protein Simple) detection. Normalized chemiluminescence intensity was automatically calculated by Compass Simple Western software (Version 6.1.0, Protein Simple).

### Statistical analysis of phenotypic data

Statistical analysis was conducted using R Version 4.4.1, with statistical significance set at *p < .05*. Linear and logistic regression models were used to examine associations between CNV status (control, deletion, duplication) and dimensional psychiatric/cognitive outcomes and binary diagnostic outcomes, respectively, adjusting for age and sex. Detailed statistical methodology is provided in **Supplementary materials and methods.**

### Statistical analysis of biomolecular data

Statistical analysis was conducted using R Version 4.4.1, with statistical significance set at *p < .05*. Protein levels were compared across control, deletion and duplication groups using one-way ANOVA or Kruskall-Wallis test, and associations with CNV status were examined using linear regression adjusted for age and sex. Detailed statistical methodology is provided in **Supplementary materials and methods.**

Based on the results of the previous analyses, causal mediation analysis was also conducted to examine whether protein levels mediated the association between CNV status and autism symptom count (see **Supplementary materials and methods).**

## Results

### Phenotypic assessments

See **Supplementary Table 2** for rates of diagnoses for each psychiatric outcome, mean symptom counts, and mean IQ scores.

#### Comparison with sibling controls confirms psychiatric outcomes associated with 16p11.2 CNV syndromes in CYP with deletions or duplications

CYP with 16p11.2 deletion and duplication syndromes showed increased ADHD symptoms compared with controls, and higher ADHD symptoms in duplication relative to deletion (**Figure 1 and Supplementary Table 3)**. Both 16p11.2 deletion and duplication groups exhibited elevated anxiety symptoms, mood disorder symptoms, and autistic traits compared with controls, with no differences between deletion and duplication. Elevated ODD symptoms were observed only in CYP with 16p11.2 duplication, who also showed higher ODD symptoms than controls and those with deletion. DCDQ scores were inverted for ease of interpretation, such that higher scores indicated poorer motor function; both CYP with deletion and duplication showed greater motor coordination difficulties than controls, with no differences between deletion and duplication. OCD symptoms were comparable between CYP with 16p11.2 deletion or duplication and controls.

**Figure 1.**
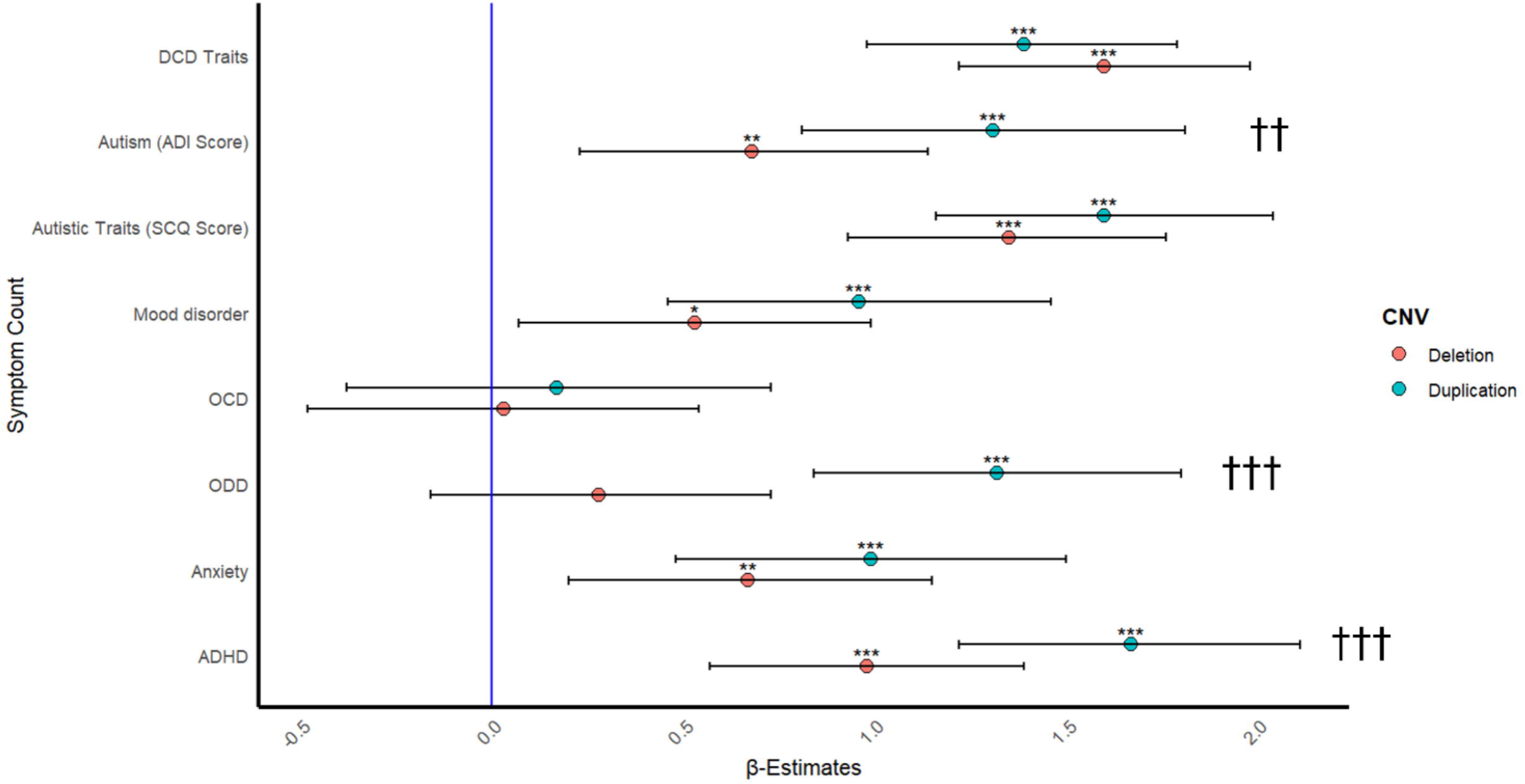
Forest plot showing beta estimates (β) and 95% confidence intervals (CIs) for psychiatric symptom counts in 16p11.2 deletion and duplication carriers relative to controls. Each point represents the estimated regression coefficient (β) for a given psychiatric outcome in CYP with 16p11.2 deletion and duplication compared with controls. Error bars indicate 95% confidence intervals. The vertical reference line at β = 0 represents no difference from controls; positive coefficients indicate higher symptom counts relative to controls. Asterisks denote statistically significant differences from controls (\**p* < .05, **p < .01, \*\*\**p* < .001). Daggers denote statistically significant differences between deletion and duplication groups (†*p* < .05, ††*p* < .01, †††*p* < .001).

CYP with 16p11.2 deletion and duplication exhibit higher autism symptoms than controls across all autism subdomains (**Supplementary Table 5)**. CYP with 16p11.2 duplication exhibit increased autism symptoms compared to those with deletion overall (ADI-R total score) and on the subdomains of communication and language (ADI-R subdomain B) and RRSB (ADI-R subdomain C).

See also **Supplementary Table 4** for additional analyses relating to binary psychiatric diagnoses and cognitive outcomes.

#### Reduced cognitive ability in CYP with 16p11.2 deletion and duplication syndromes compared to sibling controls

CYP with deletion and duplication have lower FSIQ (−21 and −23 IQ points respectively), VIQ (−21 and −19 IQ points respectively) and PIQ (−18 and −29 IQ points respectively), compared to controls. There is no difference between CYP with deletion and duplication across any of the IQ categories **(Figure 2).**

**Figure 2.**
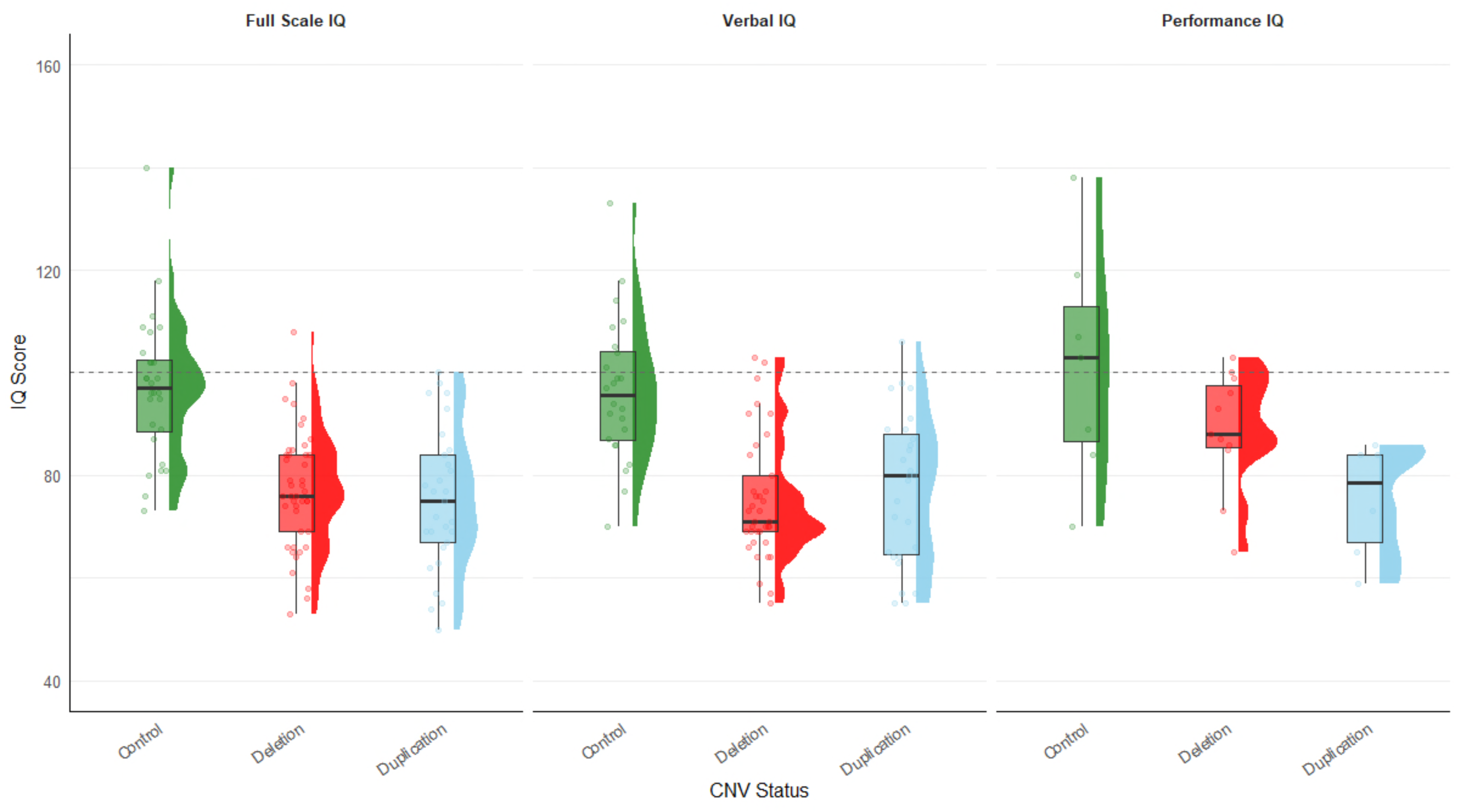
Distribution of Full-Scale IQ (FSIQ), Verbal IQ (VIQ), and Performance IQ (PIQ) scores. Each panel includes a box plot with a scatter overlay and a half violin plot to the side, providing a comprehensive view of the data distribution for each IQ measure. The central line indicates the median, and the box spans the interquartile range (IQR). The whiskers extend to 1.5 times the IQR. Individual data points are plotted over the box plot to show the distribution and density of the data. Points are jittered for clarity. The half violin plot shows the density distribution of the data. Each panel compares the IQ scores of controls and those with 16p11.2 deletion and duplication. The left panel illustrates Full Scale IQ, the central panel verbal IQ, and the right panel, performance IQ. **Note.** Sample sizes for PIQ are smaller than for FSIQ and VIQ because PIQ data were unavailable for participants assessed remotely.

To assess potential participation bias, a binary indicator of successful blood sample collection (yes/no) was added to each linear regression model. Successful blood sample collection was not significantly associated with any psychiatric outcome, and inclusion of this variable did not alter the pattern of results (**Supplementary Table 6**). Although successful blood sample collection was associated with higher PIQ, this finding likely reflects differences in participation and assessment context rather than a direct relationship between cognitive ability and blood taking; participants who completed in-person assessments were both more likely to provide a blood sample and to have complete PIQ data available. As PIQ scores were unavailable for participants assessed remotely, differences in assessment modality may have contributed to the observed association.

Importantly, adjustment for blood sample collection status did not alter the main psychiatric or cognitive findings, suggesting that selection bias related to blood sample provision was limited and the subsequent biomarker analyses were not substantially biased by differential sample donation.

### Biomarker analysis

#### Peripheral ERK1 and phospho-ERK1 levels show opposing changes in CYP with 16p11.2 deletion and duplication

We first analysed the expression levels of total ERK1 and ERK2, and their phosphorylated forms, in PBMCs from CYP with 16p11.2 deletion, duplication and controls. As the MAPK3 gene, coding for ERK1, is within the 16p11.2 region, we expected gene dosage-dependent changes in protein levels. In addition, we investigated potential compensatory alterations in ERK2 encoded by MAPK1, located on another chromosome.

Peripheral ERK1 (**Figure 3A-C)** and phospho-ERK1 (**Figure 3D**-**F**) are decreased in participants with 16p11.2 deletion compared to controls. In participants with 16p11.2 duplication, ERK1 and phospho-ERK1 are significantly increased compared to the deletion, with a trend toward increase compared to controls (**Supplementary Table 7**). No compensatory changes were observed for either ERK2 (**Figure 3B-C**) or phospho-ERK2 (**Figure 3E-F**) (**Supplementary Table 7**).

**Figure 3.**
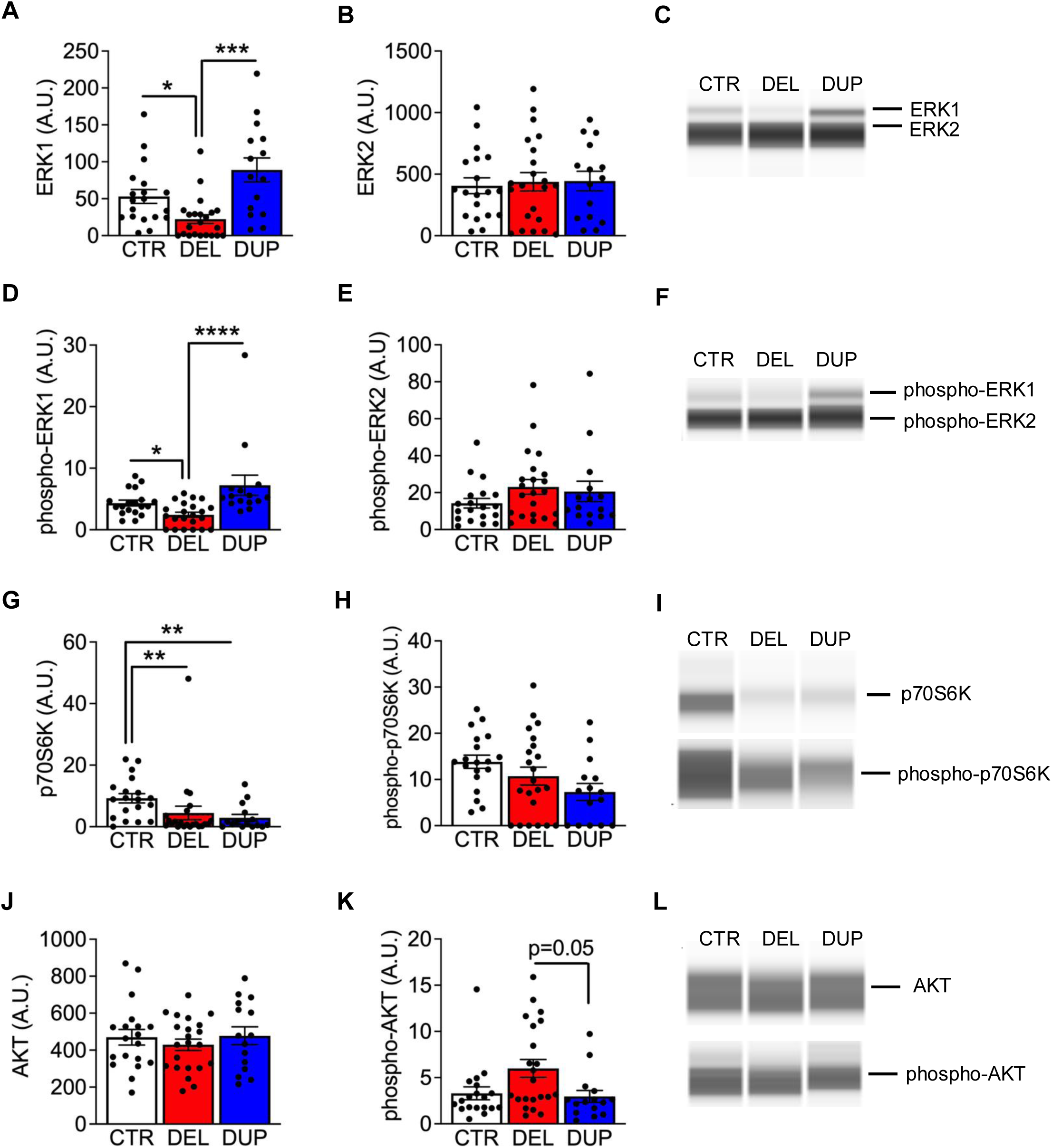
Quantification of ERK1/2, p70S6K and AKT protein levels, and their active forms, in the PBMCs of CYP with 16p11.2 deletion (DEL), duplication (DUP) and control siblings (CTR). Protein lysates from PBMCs were subjected to Automated Western blot analysis (Jess, Protein Simple). Quantification of protein levels and representative pictures are shown in A-C (total ERK1 and ERK2), D-F (phospho-ERK1 and phospho-ERK2), G-I (total and phospho-p70S6K), J-L (total and phospho-AKT). Data are shown as mean ± sem. A.U.: arbitrary units. *p<.05, **p<.01, ***p<.001, ****p<.0001.

Linear regression analyses confirmed that the deletion is predictive of both ERK1 and phospho-ERK1, but not ERK2 or phospho-ERK2 levels (**Supplementary Table 8**).

#### p70S6K is significantly downregulated in PBMCs of both CYP with 16p11.2 deletion and duplication compared to controls

As the PI3K-AKT-mTOR pathway is involved in neurodevelopmental disorders (18), we investigated potential alterations in signalling proteins along this pathway in deletion and duplication groups.

We found a downregulation of p70S6K, a downstream mediator of mTORC1 complex, in both deletion and duplication compared to controls (**Figure 3G-I**), with no differences between the deletion and the duplication groups (**Supplementary Table 7**). Phospho-p70S6K only shows a trend towards reduction in both the deletion and duplication (**Figure 3H**-**I**) (**Supplementary Table 7**).

Linear regression analysis confirmed that both deletion and duplication are predictive of p70S6K, but not phospho-p70S6K levels, with no effect of age and sex (**Supplementary Table 8**).

### Phospho-AKT levels are increased in CYP with 16p11.2 deletion compared to those with 16p11.2 duplication

We next assessed whether the observed p70S6K changes were influenced by the expression of the upstream modulators TSC1 and AKT. However, no significant alterations in either AKT (**Figure 3J**-**L**) or TSC1 (**Figure 4**) were detected (**Supplementary Table 7**). Only a trend toward increased phospho-AKT can be observed in individuals with the 16p11.2 deletion compared to controls as well as individuals with the duplication (p=0.05) (**Figure 3K-L**) (**Supplementary Table 7**).

**Figure 4.**
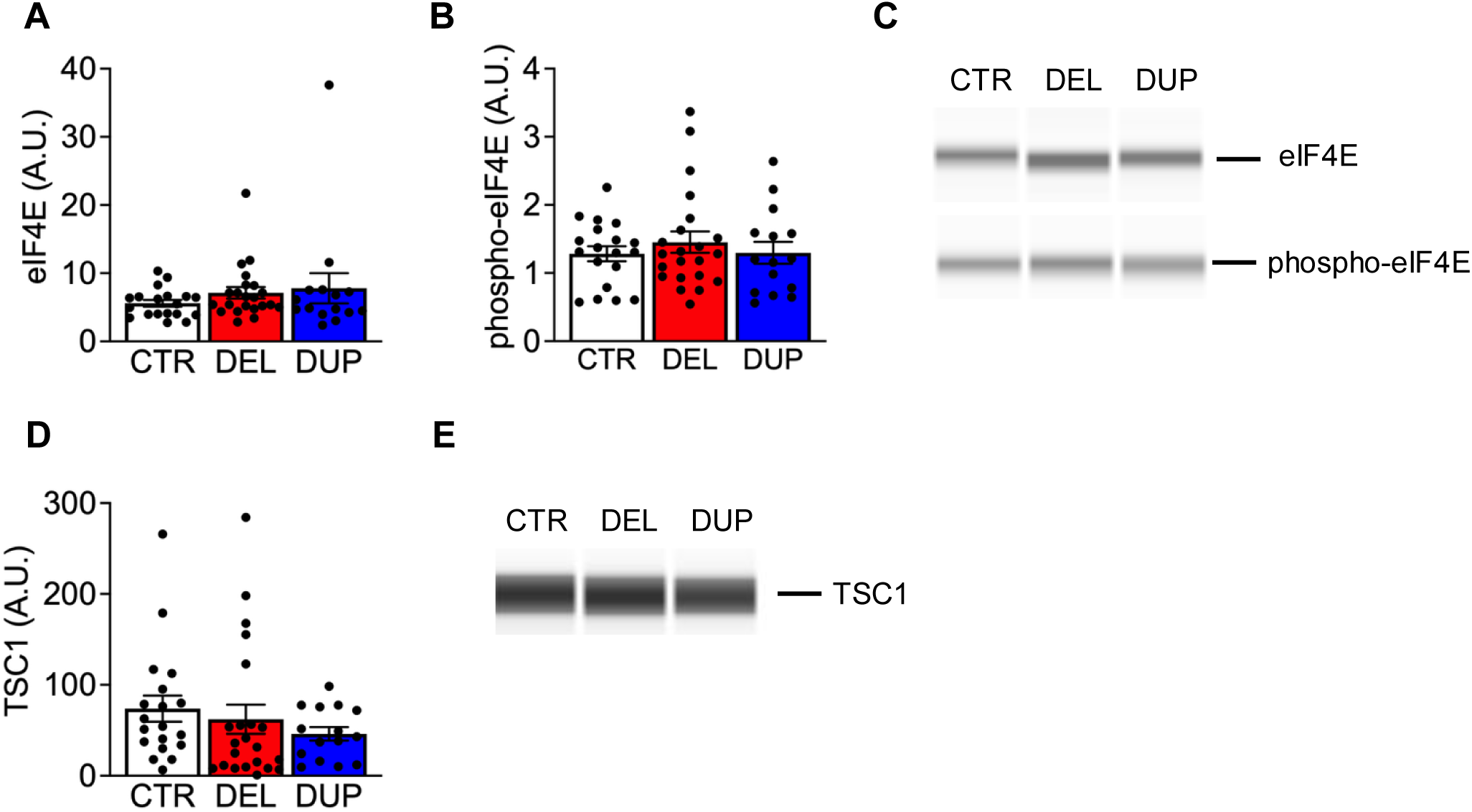
Quantification of eIF4E, and their active form, and TSC1 protein levels, in the PBMCs of CYP with 16p11.2 CNVs and control siblings. Protein lysates from PBMCs were subjected to Automated Western blot analysis (Jess, Protein Simple). Quantification of protein levels and representative pictures are shown in A-C (total and phospho-eIF4E) and D-E (TSC1). Data are shown as mean ± sem. A.U.: arbitrary units.

We next investigated whether the translation initiation factor eIF4E, downstream p70S6K, was impacted by p70S6K changes. However, both total (**Figure 4A-C**) and phospho-eIF4E (**Figure 4B** and **C**) exhibit similar expression levels across genotypes (**Supplementary Table 7**).

Subsequent linear regression analyses did not detect any relationship between these proteins and the CNV status (**Supplementary Table 8**).

Altogether, our biomarkers analysis confirms that peripheral ERK1 and phospho-ERK1 levels reflect opposing changes in MAPK3/ERK1 gene dosage in individuals with 16p11.2 CNVs, with no compensatory changes in either ERK2 or phospho-ERK2. Our data also demonstrate alterations in key components of the protein synthesis pathway, with a significant downregulation of p70S6K in both CNV groups and a slight increase in AKT activity in individuals with the 16p11.2 deletion.

#### Peripheral ERK1 and phospho-ERK1 alterations are not associated with the severity of cognitive, neurodevelopmental and psychiatric symptom severity

Based on the observed alterations in ERK1 and phospho-ERK1 levels in both CNV groups, we performed a linear regression to assess whether ERK1, or phospho-ERK1 altered levels could predict symptom severity. ERK1 protein levels neither predict ASD traits nor DCD traits. Moreover, ERK1 does not predict FSIQ score, nor symptom counts of ADHD, anxiety, mood, OCD, and ODD, or ADI-R total and ADI-R subdomains scores (**Figure 5**, **Supplementary Tables 9-10**). Similarly, no significant association was detected between phospho-ERK1 levels and symptom severity (**Figure 5**, **Supplementary Tables 11-12**).

**Figure 5.**
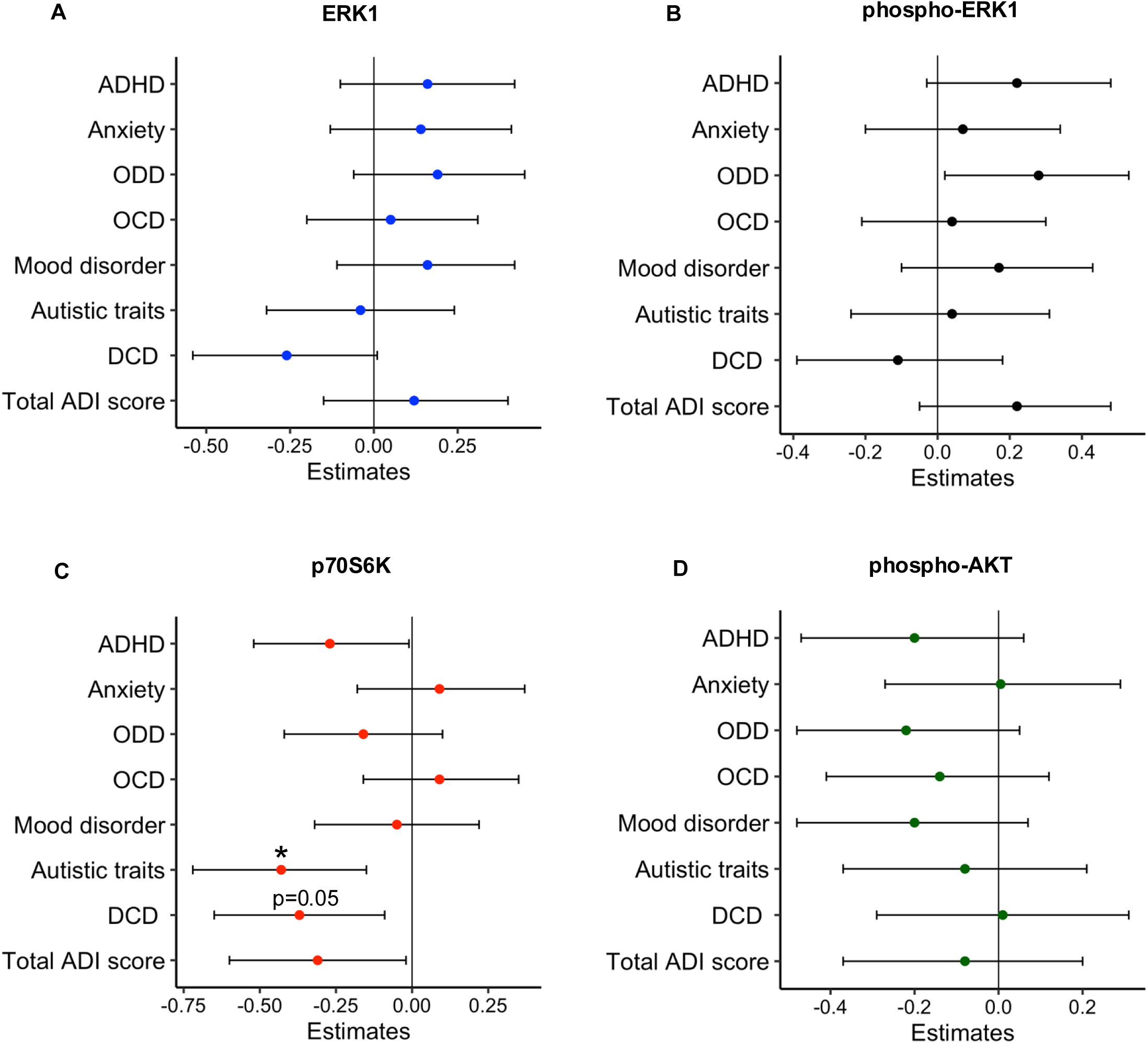
p70S6K levels are inversely associated with ASD traits. Linear regression analysis was carried out in the total sample (controls, DEL and DUP), adjusting for age and sex. Regression coefficients (β) with 95% confidence intervals indicate associations between each protein levels and symptom counts across diagnostic groups. Each point represents the estimated β for the symptom counts for a specific psychiatric measure, with horizontal error bars indicating the lower and upper confidence interval bounds. *p<.05.

#### Peripheral p70S6K protein levels are associated with the severity of ASD and DCD traits

Given the significant disruption of peripheral p70S6K observed in CYP with 16p11.2 deletion and duplication, we examined the association between p70S6K expression and symptom severity across controls and the pooled deletion and duplication sample since p70S6K levels were similar in these two groups.

We found that p70S6K is a significant predictor of ASD traits. A borderline association was detected between p70S6K and DCD traits (p_adj_=0.05), whereas no associations were found with FSIQ score, ADHD, ADI-R total, anxiety, mood, OCD and ODD (**Supplementary Table 13**). After extending our analysis to ADI-R subdomain scores, we observed that p70S6K is a significant predictor for ADI-R subdomain A and B, but not subdomain C **(Supplementary Table 14**). However, it is important to note that the ADI-R analysis was conducted only in the subsets of participants meeting criteria for ADI-R administration, which may have limited the power to detect associations.

These findings identify peripheral p70S6K levels as a potential candidate biomarker associated with ASD trait severity in the pooled 16p11.2 deletion and duplication sample. However, they do not establish that p70S6K is independently associated with ASD traits or have a causal role in their development. Adjustment for CNV genotype attenuated the association between p70S6K levels and ASD trait severity, rendering it no longer statistically significant (**Supplementary Tables 15-16**), suggesting that the observed association may be partly attributable to genotype-related differences. This should not be interpreted as definitive evidence that p70S6K has no genotype-independent association with ASD traits.

To examine whether peripheral p70S6K levels might lie on the pathway between 16p11.2 CNV status and ASD traits, we conducted a mediation analysis, which found no mediation effect of p70S6K on the 16p11.2 CNV-autism association (**Supplementary material and methods**, **Supplementary Figure 1)**.

#### Phospho-AKT levels do not correlate with cognitive, neurodevelopmental and psychiatric symptom severity

Based on the observation of increased peripheral phospho-AKT in the deletion group, we investigated the association between phospho-AKT levels and symptom severity. We found no associations with any psychiatric, cognitive or neurodevelopmental outcome (**Figure 5, Supplementary Tables 17-18**).

## Discussion

Individuals with CNVs, including 16p11.2 deletions and duplications, are at elevated risk for neurodevelopmental and mental health conditions (24), yet the molecular mechanisms linking these variants to clinical phenotypes remain unclear. Diagnosis currently relies on behavioural and neurocognitive assessments, which are costly, time-consuming, and require multiple clinical settings. Identifying reliable peripheral biomarkers could enable earlier detection of symptoms, complement existing clinical evaluations, and provide prognostic information regarding disease trajectory. PBMCs have shown promise for biomarker discovery across neurological and psychiatric conditions, including ASD (25–30), though this approach has not yet been applied to 16p11.2 CNVs. Preclinical evidence suggests that multiple genes within the 16p11.2 locus converge on shared signalling pathways, with MAPK3 emerging as a key regulator of the mTOR cascade—implicated in ASD, ADHD, and cognitive dysfunction (21, 31). Recent evidence demonstrated mTOR signalling dysregulation in neural precursor cells (NPCs) from individuals with idiopathic autism and 16p11.2 deletion, leading to deficits in neurite outgrowth and cell migration, further supporting the role of protein translation control in neurodevelopmental disorders (32).

As detecting signalling changes in the brain is not currently feasible, it is imperative to determine whether molecular alterations can be also detected peripherally. The current study is unique because it combines a deep characterisation of the neurodevelopmental, psychiatric, and cognitive profiles of individuals with 16p11.2 deletions and duplications, compared to controls, with the profiling of key signalling proteins in PBMCs, focusing on the ERK and mTOR pathways, to evaluate pathway-specific alterations and their associations with clinical outcomes.

In line with previous research (3), CYP with 16p11.2 deletions or duplications in our sample had significantly lower FSIQ, VIQ and PIQ compared to their siblings without neurodevelopmental risk CNV. Additionally, CYP with 16p11.2 deletion or duplication were more likely to have increased symptom counts for anxiety, mood disorder, and ASD, as well as lower cognitive abilities, reflecting the well-established risk profile associated with the 16p11.2 locus (3). We also revealed that individuals with 16p11.2 duplication are more likely to have increased symptom counts for ODD, and ADHD, than young people with the deletion. Individuals with 16p11.2 duplication were also more likely than those with the deletion to have increased autism symptom counts on the ADI-R, and specifically, on sub-domain B; language and communication. These differing profiles between CYP with deletion and duplication, with distinct patterns emerging across specific diagnostic domains, suggest that reciprocal CNVs at 16p11.2 may exert both common and unique effects on neurodevelopmental outcome. This highlights the importance of detailed and wide-ranging phenotyping to allow detection of specific risk profiles in individuals with these high-risk genomic variants.

Our proteomic results revealed genotype-specific alterations in peripheral ERK1 (MAPK3) and its phosphorylated form, with upregulation in 16p11.2 duplication and downregulation in the deletion, confirming reliable detection of CNV-related changes in PBMCs. This pattern is consistent with the opposing neuroanatomical phenotypes reported in individuals with 16p11.2 CNVs (33). However, other signalling proteins did not display these reciprocal changes. For instance, both CNV groups showed downregulation of p70S6K, while phospho-AKT was increased in individuals with the deletion compared to the duplication, implicating mTOR pathway disruption. While ERK, p70S6K, and AKT all converge on mTOR, their differential expression patterns suggest that mTOR dysregulation cannot be explained solely by gene dosage, pointing to more complex regulatory mechanisms. Our findings of disrupted mTOR signalling in this group at genetic risk of NDDs align with evidence which demonstrated that dysfunctional mTOR signalling alters spine protein translation, thereby determining changes in spine morphology and density, features that have been found in individuals with ASD (34). These findings are also in line with a recent proteomic study on induced pluripotent stem cell (iPSC)-derived neural progenitors from individuals with idiopathic ASD and 16p11.2 deletion, which identified two cell populations showing up- and downregulation of mTOR, both displaying similar defects in neurite outgrowth and cell migration, despite genetic heterogeneity (32).

Notably, we found that p70S6K levels predict the severity of ASD traits, a core symptom of 16p11.2 CNVs. This finding gives initial evidence by association that p70S6K could be a candidate peripheral biomarker for neurodevelopmental symptom burden in individuals with 16p11.2 CNVs, potentially reflecting underlying disruptions in mTOR pathway signalling (but see limitations below). In contrast, AKT activity, despite being increased in 16p11.2 deletion, does not correlate with the severity of any neuropsychiatric, cognitive and neurodevelopmental symptoms. These findings align with previous post-mortem studies of idiopathic ASD, which reported reductions in multiple mTOR-related proteins, including p70S6K, in cortical tissue (35). These converging lines of evidence reinforce the hypothesis that mTOR dysregulation, and specifically altered p70S6K activity, may contribute to atypical neurodevelopment across both genetically at-risk, and idiopathic presentations. Given p70S6K’s role in regulating protein synthesis and synaptic plasticity, its association with symptom severity underscores the translational relevance of peripheral proteomic profiling in stratifying risk and guiding future therapeutic targets.

Finally, we have considered some technical limitations of our study. Firstly, genotype-first recruitment may favour more severely affected individuals referred for genetic testing, although this cohort reflects those most likely to require clinical support. Secondly, blood-sample participation may also have introduced selection bias, but measured outcomes did not differ significantly between those who did and did not provide samples. Thirdly, the targeted analysis enabled hypothesis-driven investigation of ERK and mTOR signalling but may have missed broader proteomic changes, and peripheral protein levels may not directly reflect brain processes. Furthermore, the cross-sectional design and young cohort preclude conclusions about causality, developmental trajectories and later-emerging psychiatric outcomes. Finally, the modest sample size limited our genotype-adjusted and mediation analyses. Therefore, the absence of significant mediation by p70S6K does not exclude a small effect. Despite these limitations, our findings give novel support for p70S6K as a candidate peripheral correlate of ASD trait severity in individuals with 16p11.2 deletion and duplication. These findings provide a strong logic for further larger and longitudinal studies with unbiased proteomic approaches and replication to further consolidate its role as a true biomarker.

## Supporting information

https://github.com/CardiffMRCPathfinder/NeurodevelopmentalCNVCalling

## Acknowledgements

We would like to thank all the children and their family members who took part in this study, conducted during Covid-19 pandemic, as well as all the support we have had from NHS medical genetic clinics, and support charities, including Unique. This research was funded by the Medical Research Council as part of the ‘Therapeutic target validation in mental health’ call (MR/S037667/1 to RB, JG and MvdB). MvdB received additional support from the Medical Research Council (MR/T033045/1, MR/W014416/1, MR/W028395/1, MR/W020297/1, MR/V004905/1RE), Wellcome Trust (227882/Z/23/Z, 226709/Z/22/Z) and National Institutes of Mental Health (U01MH119758) and RB was supported by the NextGeneration EU (NGEU), the Ministry of University and Research (MUR), National Recovery and Resilience Plan (NRRP), project MNESYS (PE0000006) – A multiscale integrated approach to the study of the nervous system in health and disease (DN. 1553 11 October 2022). This work is dedicated to the memory of Dr. Kishan Sharma, who was involved in its early stages.

## References

1. Mannik K, Magi R, Mace A, Cole B, Guyatt AL, Shihab HA, et al. Copy number variations and cognitive phenotypes in unselected populations. JAMA. 2015;313(20):2044–54.

2. Cunningham AC, Hall J, Einfeld S, Owen MJ, consortium I-I, van den Bree MBM. Assessment of emotions and behaviour by the Developmental Behaviour Checklist in young people with neurodevelopmental CNVs. Psychol Med. 2022;52(3):574–86.

3. Niarchou M, Chawner S, Doherty JL, Maillard AM, Jacquemont S, Chung WK, et al. Psychiatric disorders in children with 16p11.2 deletion and duplication. Transl Psychiatry. 2019;9(1):8.

4. D’Angelo D, Lebon S, Chen Q, Martin-Brevet S, Snyder LG, Hippolyte L, et al. Defining the Effect of the 16p11.2 Duplication on Cognition, Behavior, and Medical Comorbidities. JAMA Psychiatry. 2016;73(1):20–30.

5. Cunningham AC, Hall J, Owen MJ, van den Bree MBM. Coordination difficulties, IQ and psychopathology in children with high-risk copy number variants. Psychol Med. 2021;51(2):290–9.

6. Ali NMH, Chawner S, Kushan-Wells L, Bearden CE, Mulle JG, Pollak RM, et al. Comparison of autism domains across thirty rare variant genotypes. EBioMedicine. 2025;112:105521.

7. Chawner S, Doherty JL, Anney RJL, Antshel KM, Bearden CE, Bernier R, et al. A Genetics-First Approach to Dissecting the Heterogeneity of Autism: Phenotypic Comparison of Autism Risk Copy Number Variants. Am J Psychiatry. 2021;178(1):77–86.

8. Chawner S, Owen MJ, Holmans P, Raymond FL, Skuse D, Hall J, et al. Genotype-phenotype associations in children with copy number variants associated with high neuropsychiatric risk in the UK (IMAGINE-ID): a case-control cohort study. Lancet Psychiatry. 2019;6(6):493–505.

9. Moufawad El Achkar C, Rosen A, Kessler SK, Steinman KJ, Spence SJ, Ramocki M, et al. Clinical Characteristics of Seizures and Epilepsy in Individuals With Recurrent Deletions and Duplications in the 16p11.2 Region. Neurol Genet. 2022;8(5):e200018.

10. Gur R, Bearden C, Jacquemont S, Jizi K, Amelsvoort van T, van den Bree M, et al. Neurocognitive Profiles of 22q11.2 and 16p11.2 Deletions and Duplications. Res Sq. 2023.

11. Leone R, Zuglian C, Brambilla R, Morella I. Understanding copy number variations through their genes: a molecular view on 16p11.2 deletion and duplication syndromes. Front Pharmacol. 2024;15:1407865.

12. Auwerx C, Kutalik Z, Reymond A. The pleiotropic spectrum of proximal 16p11.2 CNVs. Am J Hum Genet. 2024;111(11):2309–46.

13. Rein B, Yan Z. 16p11.2 Copy Number Variations and Neurodevelopmental Disorders. Trends Neurosci. 2020;43(11):886–901.

14. More L, Lauterborn JC, Papaleo F, Brambilla R. Enhancing cognition through pharmacological and environmental interventions: Examples from preclinical models of neurodevelopmental disorders. Neurosci Biobehav Rev. 2020;110:28–45.

15. Indrigo M, Morella I, Orellana D, d’Isa R, Papale A, Parra R, et al. Nuclear ERK1/2 signaling potentiation enhances neuroprotection and cognition via Importinalpha1/KPNA2. EMBO Mol Med. 2023;15(11):e15984.

16. Mendoza MC, Er EE, Blenis J. The Ras-ERK and PI3K-mTOR pathways: cross-talk and compensation. Trends Biochem Sci. 2011;36(6):320–8.

17. Iroegbu JD, Ijomone OK, Femi-Akinlosotu OM, Ijomone OM. ERK/MAPK signalling in the developing brain: Perturbations and consequences. Neurosci Biobehav Rev. 2021;131:792–805.

18. Borrie SC, Brems H, Legius E, Bagni C. Cognitive Dysfunctions in Intellectual Disabilities: The Contributions of the Ras-MAPK and PI3K-AKT-mTOR Pathways. Annu Rev Genomics Hum Genet. 2017;18:115–42.

19. Pucilowska J, Vithayathil J, Pagani M, Kelly C, Karlo JC, Robol C, et al. Pharmacological Inhibition of ERK Signaling Rescues Pathophysiology and Behavioral Phenotype Associated with 16p11.2 Chromosomal Deletion in Mice. J Neurosci. 2018;38(30):6640–52.

20. Pucilowska J, Vithayathil J, Tavares EJ, Kelly C, Karlo JC, Landreth GE. The 16p11.2 deletion mouse model of autism exhibits altered cortical progenitor proliferation and brain cytoarchitecture linked to the ERK MAPK pathway. J Neurosci. 2015;35(7):3190–200.

21. Lee DY. Roles of mTOR Signaling in Brain Development. Exp Neurobiol. 2015;24(3):177–85.

22. Eaves LC, Wingert HD, Ho HH, Mickelson EC. Screening for autism spectrum disorders with the social communication questionnaire. J Dev Behav Pediatr. 2006;27(2 Suppl):S95–S103.

23. Lord C, Rutter M, Le Couteur A. Autism Diagnostic Interview-Revised: a revised version of a diagnostic interview for caregivers of individuals with possible pervasive developmental disorders. J Autism Dev Disord. 1994;24(5):659–85.

24. Coe BP, Witherspoon K, Rosenfeld JA, van Bon BW, Vulto-van Silfhout AT, Bosco P, et al. Refining analyses of copy number variation identifies specific genes associated with developmental delay. Nat Genet. 2014;46(10):1063–71.

25. Gladkevich A, Kauffman HF, Korf J. Lymphocytes as a neural probe: potential for studying psychiatric disorders. Prog Neuropsychopharmacol Biol Psychiatry. 2004;28(3):559–76.

26. Nishimura Y, Martin CL, Vazquez-Lopez A, Spence SJ, Alvarez-Retuerto AI, Sigman M, et al. Genome-wide expression profiling of lymphoblastoid cell lines distinguishes different forms of autism and reveals shared pathways. Hum Mol Genet. 2007;16(14):1682–98.

27. Hu VW, Nguyen A, Kim KS, Steinberg ME, Sarachana T, Scully MA, et al. Gene expression profiling of lymphoblasts from autistic and nonaffected sib pairs: altered pathways in neuronal development and steroid biosynthesis. PLoS One. 2009;4(6):e5775.

28. Rosina E, Battan B, Siracusano M, Di Criscio L, Hollis F, Pacini L, et al. Disruption of mTOR and MAPK pathways correlates with severity in idiopathic autism. Transl Psychiatry. 2019;9(1):50.

29. Gazestani VH, Pramparo T, Nalabolu S, Kellman BP, Murray S, Lopez L, et al. A perturbed gene network containing PI3K-AKT, RAS-ERK and WNT-beta-catenin pathways in leukocytes is linked to ASD genetics and symptom severity. Nat Neurosci. 2019;22(10):1624–34.

30. Pagano J, Perez Arevalo A, Nosanova A, Bauer HF, Loth E, Andres S, et al. SHANK3 and beta-synuclein are novel blood-based biomarkers for the Phelan-McDermid Syndrome: a pilot study. Transl Psychiatry. 2026;16(1).

31. Kumar RA, Marshall CR, Badner JA, Babatz TD, Mukamel Z, Aldinger KA, et al. Association and mutation analyses of 16p11.2 autism candidate genes. PLoS One. 2009;4(2):e4582.

32. Prem S, Dev B, Peng C, Mehta M, Alibutud R, Connacher RJ, et al. Dysregulation of mTOR signaling mediates common neurite and migration defects in both idiopathic and 16p11.2 deletion autism neural precursor cells. Elife. 2024;13.

33. Martin-Brevet S, Rodriguez-Herreros B, Nielsen JA, Moreau C, Modenato C, Maillard AM, et al. Quantifying the Effects of 16p11.2 Copy Number Variants on Brain Structure: A Multisite Genetic-First Study. Biol Psychiatry. 2018;84(4):253–64.

34. Hutsler JJ, Zhang H. Increased dendritic spine densities on cortical projection neurons in autism spectrum disorders. Brain Res. 2010;1309:83–94.

35. Nicolini C, Ahn Y, Michalski B, Rho JM, Fahnestock M. Decreased mTOR signaling pathway in human idiopathic autism and in rats exposed to valproic acid. Acta Neuropathol Commun. 2015;3:3.

