## Supplementary material for "p70 ribosomal protein S6 Kinase (p70S6K) as a potential peripheral biomarker for autistic symptoms in 16p11.2 deletion and duplication syndromes": https://github.com/CardiffMRCPathfinder/NeurodevelopmentalCNVCalling

### Supplementary Materials and Methods

#### Genotyping

We screened for the presence or absence of 54 CNVs considered to be recurrent, pathogenic, or likely pathogenic, as per the American College of Medical Genetics and Genomics guidelines (1) and associated with high risk of NDD (ND-CNV) (2). We confirmed the presence of a proximal 16p11.2 deletion or duplication in diagnosed participants and the absence in controls, using the Cardiff University Centre for Neuropsychiatric Genetics and Genomics (CNGG) DRAGON pipeline. Briefly, we performed initial QC excluding samples based on 100 or more CNVs, waviness factor ≤0.037 or >0.037, or log R ratio SD >0.24. We visually inspected log R ratios and B-allele frequencies to exclude low-quality calls. This pipeline is described in more detail in (3) (see also: https://github.com/CardiffMRCPathfinder/NeurodevelopmentalCNVCalling) and has been used across a range of studies (e.g.,(2, 4)).

Participants with 16p11.2 CNV were excluded if we found they had a second hit of any of the other ND-CNV we screened for. Participants were excluded as controls if they were found to have a ND-CNV and were recategorised into the deletion/duplication groups where relevant.

#### Statistical analysis of symptom counts and IQ scores

Linear regression analyses were fitted using the lm() function for each psychiatric symptom count (ADHD, Anxiety, ODD, OCD, mood disorder, autistic traits (SCQ score) autism (ADI-R total score) and DCD traits) as well as IQ scores (FSIQ, PIQ and VIQ), with CNV status (control, deletion, duplication) as a three-level factor, whilst controlling for age and sex. Dimensional symptom scores were transformed using the Tukey Ladder of Powers to ensure that the data fitted the normal distribution as closely as possible. Transformed scores were then standardized into z-scores to facilitate interpretation across variables. Beta estimates were estimated for deletion versus control and duplication versus control. Secondary analyses using post-hoc contrasts comparing deletion and duplication carriers were derived using estimated marginal means.

Primary hypotheses tested the association of deletion versus control and duplication versus control status with each psychiatric/cognitive outcome (10 outcomes). Due to zero instances of OCD and ID in the control group, regression models for OCD and ID did not converge and valid p-values could not be obtained. OCD and ID were therefore excluded from multiple-comparison correction procedures. For each of the eight models that yielded valid p-values, two planned comparisons were conducted (deletion vs control and duplication vs control), yielding a total of 16 hypothesis tests. To control the false discovery rate across these multiple comparisons, p-values were adjusted using the Benjamini–Hochberg (BH) procedure. The BH correction was applied across all contrasts jointly, and unadjusted (p), and adjusted ($p_{adj}$) p-values are reported. Comparisons between deletion and duplication carriers addressed a separate hypothesis and were corrected independently across outcomes (10 outcomes); again, due to zero instances of OCD in the duplication group, valid p-values could not be obtained, and thus OCD was excluded from the multiple comparison corrections.

Where SCQ data was available to assess autistic traits (n = 26 control, 41 deletion, 26 duplication), linear regression models were fitted to investigate associations between CNV status (deletion, duplication, control), and autistic symptoms using ADI-R total and subdomain scores (Social Interaction, Communication, RRSB), whilst controlling for age and sex. Where individuals scored <15 on the SCQ, we coded ADI scores as 0. Dimensional symptom scores were transformed via the Tukey Ladder of Powers and standardized into z-scores across the subgroup. The BH corrections were applied, correcting for the autism symptom domains evaluated (4 outcomes) across two comparisons (deletion vs control and duplication vs control) (8 hypothesis tests). Post-hoc contrasts comparing deletion and duplication carriers were derived using estimated marginal means, and BH corrections were applied separately for these four tests.

Sensitivity analyses for each regression model were conducted, including “successful blood sample (yes/no)” as a covariate in all models to account for potential bias, ensuring there were no systematic differences in phenotypic outcomes between individuals who did or did not provide a blood sample, as only a subset of individuals (n = 18 controls, 23 deletion, 15 duplication) were able to provide a blood sample. In the original models 2-subtest and 4-subtest WASIs were treated equivalently; sensitivity analyses were conducted excluding 2-subtest observations.

**Statistical analysis of binary psychiatric and cognitive outcomes**

Logistic regression analyses were fitted using the glm() function for each binary psychiatric/cognitive outcome (ADHD, Anxiety, ODD, OCD, mood disorder, ASD, any psychiatric disorder, indicative DCD and ID (FSIQ≤70)), with CNV status (control, deletion, duplication) as a three-level factor, whilst controlling for age and sex. Odds ratios were estimated for deletion versus control and duplication versus control. Post-hoc contrasts comparing deletion and duplication carriers were derived using estimated marginal means. Primary hypotheses tested the association of deletion versus control and duplication versus control status with each outcome (14 tests). P-values for these comparisons were corrected using the Benjamini–Hochberg false discovery rate; due to zero instances of OCD and ID in the control group, regression models for OCD and ID did not converge and valid p-values could not be obtained. OCD and ID were therefore excluded from multiple-comparison correction procedures. Comparisons between deletion and duplication carriers addressed a separate, secondary question and were corrected independently across outcomes (8 tests); again, due to zero instances of OCD in the duplication group, valid p-values could not be obtained, and thus OCD was excluded from the multiple comparison corrections.

**Statistical analysis of biomolecular data and association analysis**

After assessing normality of data by Shapiro-Wilk test, protein levels in samples from controls, deletion and duplication groups were compared using one-way ANOVA or Kruskall-Wallis test. Isolated missing values for the protein, due to insufficient biological samples, were imputed using the mean protein value of the corresponding genotype. Where significant main effects were identified, post-hoc comparisons were performed. Linear regression analyses were performed for each protein level using the lm() function, with CNV status (control, deletion, duplication) as a three-level factor, whilst controlling for age and sex. Protein levels which did not follow a normal distribution were transformed using Tukey Ladder of Powers transformation before undergoing linear regressions. A BH correction was applied across all contrasts jointly. Beta estimates were estimated for deletion versus control and duplication versus control. As a secondary post hoc analysis, deletion group was directly compared with duplication group using estimated marginal means.

The association between protein levels and the dimensional symptom count scores (ADHD, anxiety, mood disorder, OCD, ODD, autistic traits (SCQ score), DCD traits, ADI-R, as well as FSIQ) was examined by linear regression analyses, after applying Tukey Ladder of Powers transformation on the protein levels and the raw scores for symptom counts. All transformed data were standardised into z-scores calculated across the whole sample. BH correction was applied across all contrasts jointly. For each protein, linear regression analysis was carried out with symptom scores as the dependent variable and protein levels as the predictor, adjusting for age and sex. For p70S6K, a second linear regression was carried out including also the CNV status, as a two-levels factor (control, CNV group).
Where SCQ data were available, linear regression analyses were performed to examine the associations between protein levels and ADI-R total and subdomain scores (Social Interaction, Communication, RRSB), as described in the section relative to statistical analysis of phenotypic data.

**Exploratory causal mediation analysis**

Analyses were performed using the *mediation* package in R. Two regression models were specified: (1) a mediator model estimating the effect of CNV status on protein levels, and (2) an outcome model estimating the effect of CNV status and protein levels on symptom counts. CNV status was included as the independent variable, protein level as the mediator, and symptom count as the outcome. Age and sex were included as covariates in both models. In line with the causal steps approach proposed by Baron and Kenny, mediation analyses were restricted to cases where there was evidence of significant associations between (i) CNV status and symptom count, (ii) CNV status and protein levels, and (iii) protein levels and symptom count.

The average causal mediation effect (ACME; indirect effect), average direct effect (ADE), and total effect were estimated. Statistical inference was based on quasi-Bayesian Monte Carlo simulation with 1,000 simulations to derive 95% confidence intervals and p-values. The proportion of the total effect mediated by protein levels was also calculated.

**Supplementary appendix 1**

Mediation analysis was conducted for p70S6K only as the assumptions of the Baron and Kenny causal steps framework were met specifically for the association between p70S6K and autism symptoms. Causal mediation analysis showed that CNV status was significantly associated with autism symptom count (total effect: $\beta = 0.83$, $p = 0.004$). However, there was no evidence that this association was mediated by p70S6K levels (indirect effect: $\beta = 0.11$, $p = 0.174$). The direct effect of CNV on autism symptom counts remained significant ($\beta = 0.72$, $p = 0.018$), indicating that the association is largely independent of the proposed protein mediator. The proportion of the total effect mediated by p70S6K was small and non-significant (12%, $p = 0.178$), suggesting little evidence that p70S6K meaningfully mediates the 16p11.2 deletion/duplication-autism association. See **Supplementary Figure 1** for a visualisation of these results.

**Supplementary Figure 1:**

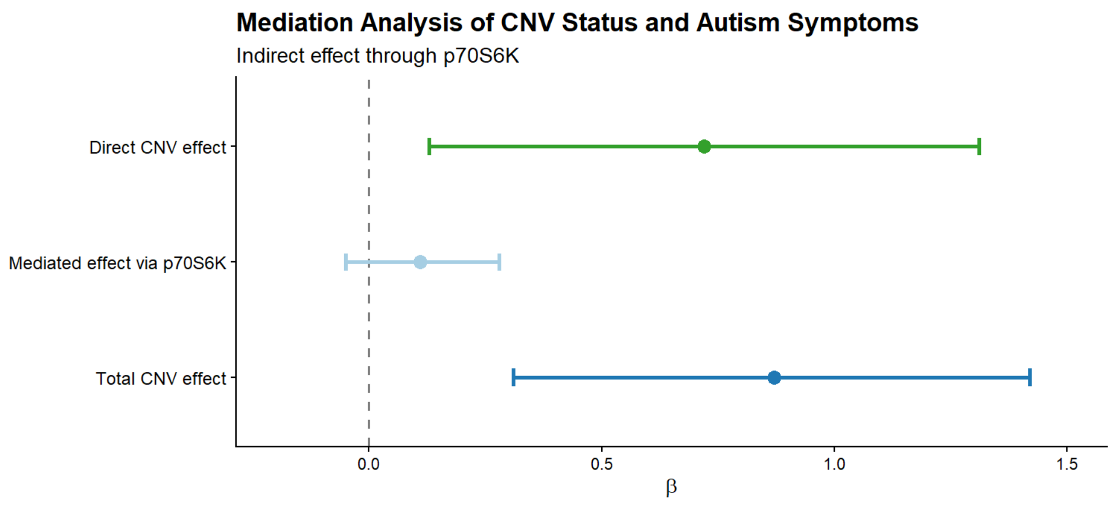

**Supplementary Table 1.** Automated Western blotting parameters and primary antibodies' details.

| Target protein | Antibody | Host species | Dilution | Blocking reagent | Stacking loading time | Sample loading time | Separation time |
|---|---|---|---|---|---|---|---|
| ERK1/2 | #4696, Cell Signalling | mouse | 1:50 | Milk-free antibody diluent | 16 sec | 6 sec | 32 min |
| Phospho-ERK1/2 | #9101, Cell Signalling | rabbit | 1:250 | Milk-free antibody diluent | 16 sec | 6 sec | 32 min |
| p70S6K | #9202, Cell Signalling | rabbit | 1:1000 | Antibody diluent 2 | 15 sec | 9 sec | 25 min |
| Phospho-p70S6K | #9206, Cell Signalling | mouse | 1:50 | Antibody diluent 2 | 18 sec | 10.8 sec | 25 min |
| AKT | #4691, Cell Signalling | rabbit | 1:10 | Antibody Diluent 2 | 15 sec | 9 sec | 25 min |
| Phospho-AKT | #4058, Cell Signalling | rabbit | 1:50 | Milk-free Diluent | 15 sec | 9 sec | 25 min |
| eIF4E | #sc27148, Santa Cruz Biotechnology | mouse | 1:50 | Antibody Diluent 2 | 18 sec | 10.8 sec | 32 min |
| Phospho-eIF4E | #9741, Cell Signalling | rabbit | 1:10 | Antibody Diluent 2 | 15 sec | 9 sec | 25 min |
| TSC1 | #6935, Cell Signalling | rabbit | 1:50 | Antibody Diluent 2 | 15 sec | 9 sec | 25 min |

**Supplementary Table 2**. Rates of diagnoses for each psychiatric outcome, symptom counts and IQ scores.

| | Deletion | Duplication | Control |
|---|---|---|---|
| **% Meeting Diagnostic Criteria** | **n = 45** | **n = 29** | **n = 30** |
| ADHD | 16 (35.5%) | 19 (65.5%) | 3 (10%) |
| Anxiety | 12 (28.9%) | 11 (37.9%) | 6 (20%) |
| ODD | 2 (4.4%) | 12 (41.4%) | 1 (3.3%) |
| OCD | 1 (2.2%) | 0 (0%) | 0 (0%) |
| Mood disorder | 2 (4.4%) | 1 (3.4%) | 1 (3.3%) |
| Any psychiatric disorder | 32 (71.1%) | 27 (93.1%) | 9 (30%) |
| | **n = 41** | **n = 27** | **n = 28** |
| Autism Spectrum Disorder (ASD) (SCQ Score ≥15) | 23 (56.1%) | 20 (71.4%) | 3 (11.1%) |
| | **n = 42** | **n = 28** | **n = 27** |
| Indicative DCD | 41 (97.6%) | 26 (92.9%) | 2 (22.2%) |
| Intellectual disability (ID = FSIQ ≤70) | 33 (26.6%) | 12 (41.4%) | 0 (0%) |
| **Mean Symptom Counts (Tukey tranformed and Z scored)** | **n = 45** | **n = 29** | **n = 30** |
| ADHD | 0.07(±0.8) | 0.75(±0.8) | -0.83 (±0.8) |
| Anxiety | 0.02(±0.9) | 0.38(±1.0) | -0.40 (±1.0) |
| ODD | -0.22(±0.8) | 0.80(±0.9) | -0.44 (±0.9) |
| OCD | -0.05(±0.9) | 0.09(±1.0) | -0.01 (±1.0) |
| Mood disorder | -0.09 (±0.9) | 0.40 (±0.9) | -0.26 (±1.1) |
| | **n = 41** | **n = 27** | **n = 28** |
| Autistic Traits: SCQ Score | 0.30 (±0.8) | 0.45 (±0.8) | -1.02 (±0.7) |
| | **n = 42** | **n = 28** | **n = 27** |
| DCD Traits: DCDQ Score | 0.53 (±0.8) | 0.29(±0.8) | -1.12 (±0.3) |
| | **n = 41** | **n = 26** | **n = 26** |
| Autistic Symptoms: (total ADI-R Score) | 0.05 (±0.92) | 0.67 (±1.13) | -0.70 (±0.34) |
| ADI-Subdomain A - Reciprocal Social Interaction | 0.05 (±1.04) | 0.55 (±1.00) | -0.70 (±0.36) |
| ADI-Subdomain B - Communication | 0.01 (±0.86) | 0.71(±1.16) | -0.70 (±0.46) |
| ADI-Subdomain C - RRSB | 0.11 (±1.01) | 0.55 (±1.08) | -0.70 (±0.30) |
| **Cognition** | **n = 37** | **n = 27** | **n = 24** |
| FSIQ | 78.08 (±11.2) | 76.56 (±13.1) | 99.04 (±14.6) |
| VIQ | 75.19 (±12.0) | 77.48 (±14.6) | 96.45 (±14.1) |
| | **n = 11** | **n = 6** | **n = 7** |
| PIQ | 88.64 (±11.6) | 75.17 (±11.3) | 101.42 (±22.8) |

**Supplementary Table 3.** Statistical results from linear regression analyses assessing the association between CNV status (deletion, duplication, control) and psychiatric symptom counts and mean IQ scores.

| Outcome | Main Effect | Deletion | Duplication | Post-Hoc |
|---|---|---|---|---|
| | | | | Deletion-Duplication |
| **ADHD** | CNV | β =0.98, CI = 0.57 - 1.39, **$p=7.10x10^{-6}$, $p_{adj}=1.42x10^{-5}$** | β =1.67, CI =1.22 - 2.11, **$p=3.99 \times 10^{-11}$, $p_{adj}=4.38 \times 10^{-10}$** | β =0.69, CI =0.29 - 1.08, **$p=7.00x10^{-4}$** |
| | Age (years) | β=0.02, CI = -0.04 - 0.08, p =0.498 | | |
| | Sex assigned at birth (females) | β =0.17, CI = -0.16 - 0.51, p =0.299 | | |
| **Anxiety** | CNV | β =0.67, CI =0.20 - 1.15, **p =0.006, $p_{adj}=7.64x10^{-3}$** | β =0.99, CI = 0.48 - 1.50, **$p=2.37x10^{-4}$, $p_{adj}=3.61x10^{-4}$** | β =0.32, CI =-0.13 - 0.76, p =0.166, p=0.332 |
| | Age (years) | β =0.09, CI =0.02 - 0.16, **p =0.009** | | |
| | Sex assigned at birth (females) | β =0.27, CI =-0.11 - 0.65, p =0.164 | | |
| **ODD** | CNV | β =0.28, CI =-0.16 - 0.73, p =0.205, $p_{adj}$=0.276 | β = 1.32, CI = 0.84 - 1.80, **$p=3.37x10^{-7}$, $p_{adj}=8.23 x10^{-7}$** | β =1.04, CI =0.621 - 1.46, **$p=3.43x10^{-6}$** |
| | Age (years) | β =0.01, CI =-0.0e6 - 0.07, p =0.785 | | |
| | Sex assigned at birth (females) | β =0.18, CI =-0.17-0.54, p =0.311 | | |
| **OCD** | CNV | β =0.03, CI =-0.48 - 0.54, p =0.907, $p_{adj}$=0.907 | β =0.17, CI = -0.38 - 0.73, p = 0.536, $p_{adj}$=0.561 | β =0.14, CI =-0.34 - 0.63, p=0.588 |
| | Age (years) | β =0.03, CI =-0.05 - 0.10, p =0.494 | | |
| | Sex assigned at birth (females) | β =0.10, CI =-0.32 - 0.51, p =0.639 | | |
| **Mood disorder** | CNV | β =0.53, CI =0.07 - 0.99, **p =0.024, $p_{adj}$=0.03** | β =0.96, CI = 0.46 - 1.46, **$p=2.37x10^{-4}$, $p_{adj}=3.61x10^{-4}$** | β =0.43, CI =-0.01 - 0.87, p=0.053 |
| | Age (years) | β =0.13, CI =0.06 - 0.19, **$p=3.79x10^{-4}$** | | |
| | Sex assigned at birth (females) | β =0.42, CI =0.04 - 0.79, **p =0.029** | | |
| **Autistic Traits (SCQ>=15)** | CNV | β =1.35, CI =0.93 - 1.76, **$p=5.06x10^{-9}$, $p_{adj}=1.85x10^{-7}$** | β =1.60, CI = 1.16 - 2.04, **$p=1.92x10^{-10}$, $p_{adj}=1.39x10^{-9}$** | β =0.25, CI =-0.13 - 0.63, p=0.199 |
| | Age (years) | β = -0.00, CI =-0.06 - 0.06, p =0.989 | | |
| | Sex assigned at birth (females) | β =0.10, CI =-0.23 - 0.43, p =0.550 | | |
| **DCD Traits** | CNV | β =1.60, CI =1.22 - 1.98, **$p=7.51x10^{-13}$, $p_{adj}=1.65x10^{-11}$** | β =1.39, CI =0.98 - 1.79, **p =1.24x10-9, $p_{adj}=6.82x10^{-9}$** | β =-0.21, CI =-0.56 - 0.141, p=0.237 |
| | Age (years) | β =-0.03, CI =-0.08 - 0.02, p =0.262 | | |
| | Sex assigned at birth (females) | β =0.03, CI =-0.27 - 0.34, p = 0.824 | | |
| **Autism (Total ADI Score)** | CNV | β =0.68, CI = 0.23 - 1.14, **p =0.004, $p_{adj}$=0.005** | β =1.31, CI =0.81 - 1.81, **$p=1.16 \times 10^{-6}$, $p_{adj}=2.64x10^{-6}$** | β =0.63, CI =0.19 - 1.06, **p=0.005** |

| | | | | |
|---|---|---|---|---|
| | Age (years) | β=-0.02, CI = -0.09 - 0.05, p =0.536 | | |
| | Sex assigned at birth (females) | β =-0.09, CI = -0.46 - 0.28, p =0.638 | | |
| **IQ DOMAINS** | | | | |
| **FSIQ** | CNV | β =-21.36, CI =-28.50 - -14.22, **p =6.21 x $10^{-8}$, $p_{adj}$=1.95x$10^{-7}$** | β =-23.13, CI = -30.75 - -15.51, **p =4.20 x $10^{-8}$, $p_{adj}$=1.84x$10^{-7}$** | β =-1.74, CI =-8.38 - 4.84, p =0.595 |
| | Age (years) | β = -0.32, CI = -1.34 - 0.71, p =0.540 | | |
| | Sex assigned at birth (females) | β =-3.36, CI =-9.15 - 2.43, p =0.252 | | |
| **VIQ** | CNV | β =-21.37, CI =-28.96 - -13.79, **p =2.63x$10^{-7}$, $p_{adj}$=7.15x$10^{-7}$** | β =-19.29, CI =-27.37 - -11.20, **p =8.64x$10^{-6}$, $p_{adj}$=1.58x$10^{-5}$** | β =2.09, CI =-4.93 - 9.10, p=0.555 |
| | Age (years) | β =0.05, CI =-1.04 - 1.14, p =0.931 | | |
| | Sex assigned at birth (females) | β =-0.83, CI =-6.98 - 5.32, p =0.789 | | |
| **PIQ** | CNV | β =-18.57, CI =-34.62 - -2.52, **p =0.026, $p_{adj}$=0.030** | β =-29.31, CI = -47.22 - -11.39, **p =0.003, $p_{adj}$=0.004** | β =-10.7, CI =-27.5 - 6.00, p=0.195 |
| | Age (years) | β =-2.29, CI =-5.02 - 0.44, p = 0.095 | | |
| | Sex assigned at birth (females) | β =-8.49, CI =-21.76 - 4.77, p =0.196 | | |

**Supplementary Table 4.** Statistical results from logistic regression analyses examining the association between CNV status (deletion, duplication and control), and binary psychiatric and cognitive outcomes.

| Outcome | Main Effect | Deletion | Duplication | Post-Hoc |
|---|---|---|---|---|
| | | | | **Deletion-Duplication** |
| **ADHD** | CNV | OR=5.37, CI=1.45 - 26.58, **p =0.020, $p_{adj}$=0.030** | OR=18.39, CI=4.65 - 97.56, **p=1.28 x $10^{-4}$, $p_{adj}$=4.48 x $10^{-4}$** | OR=3.42, CI=1.26 - 9.30, **p=0.016** |
| | Age (years) | OR=1.05, CI=0.90 - 1.24, p=0.523 | | |
| | Sex assigned at birth (females) | OR=0.97, CI=0.40 - 2.34, p=0.938 | | |
| **Anxiety** | CNV | OR=2.54, CI=0.78 - 9.21, p=0.134, $p_{adj}$=0.171 | OR=3.52, CI=1.01 - 13.53, p=0.054, $p_{adj}$=0.075 | OR=1.39, CI=0.50 - 3.89, p=0.534 |
| | Age (years) | OR=1.17, CI=0.99 - 1.39, p=0.063 | | |
| | Sex assigned at birth (females) | OR=1.48, CI=0.59 - 3.79, p=0.408 | | |
| **OCD** | CNV | Low cell counts | | Low cell counts |
| | Age (years) | OR=1.64, CI=0.62 - 5.58, p=0.312 | | |
| | Sex assigned at birth (females) | Low cell counts | | |
| **ODD** | CNV | OR=1.00, CI = 0.08 - 23.63, p =0.998, $p_{adj}$=0.998 | OR =14.28, CI =2.24 - 282.46, **p=0.018, $p_{adj}$=0.031** | OR =14.20, CI =2.76 - 73.50, p =0.002 |
| | Age (years) | OR =0.26, CI = 0.05 - 1.05, p =0.073 | | |
| | Sex assigned at birth (females) | OR =1.02, CI = 0.80 - 1.29, p =0.888 | | |
| **Mood disorder** | CNV | OR =6.74, CI = 0.43 - 230.40, p =0.205, $p_{adj}$=0.239 | OR =3.49, CI =0.10 - 136.72, p=0.455, $p_{adj}$=0.490 | OR =0.518, CI =0.032 - 8.52, p =0.645 |
| | Age (years) | OR =1.72, CI = 1.14 - 3.07, **p =0.024** | | |

| | | | | |
|---|---|---|---|---|
| | Sex assigned at birth (females) | OR =4.19, CI = 0.038 - 111.59, p =0.286 | | |
| **Autism Spectrum Disorder (SCQ>=15)** | CNV | OR =6.23, CI = 2.05 - 21.49, **p = 0.002, $p_{adj}$=0.004** | OR =12.18, CI =3.52 - 48.84, **p =1.72 x10$^{-4}$, $p_{adj}$=4.81 x10$^{-4}$** | OR = 1.95, CI = 0.69 - 5.53, p = 0.207 |
| | Age (years) | OR =0.96, CI = 0.81 - 1.13, p = 0.609 | | |
| | Sex assigned at birth (females) | OR =1.53, CI =0.63 - 3.93, p = 0.358 | | |
| **Any Psychiatric Disorder** | CNV | OR =16.71, CI = 4.58 - 77.40, **p =7.78 x10$^{-5}$, $p_{adj}$ = 3.63 x10$^{-4}$** | OR =93.55, CI =16.08 - 904.65, **p =6.46 x10$^{-6}$, $p_{adj}$=9.04 x 10$^{-5}$** | OR = 5.60, CI = 1.10 - 28.40, **p = 0.038** |
| | Age (years) | OR =1.34, CI = 1.09 - 1.71, **p =0.009** | | |
| | Sex assigned at birth (females) | OR =2.27, CI =0.79 - 7.11, p =0.140 | | |
| **Indicative DCD** | CNV | OR =83.993, CI = 13.82 - 1657.72, **p=7.01x10$^{-5}$, $p_{adj}$=3.63 x 10$^{-4}$** | OR =22.09, CI =4.86 - 162.20, **p=3.15x10$^{-4}$, $p_{adj}$=7.35x10$^{-4}$** | OR = 0.263, CI = 0.022 - 3.15, p = 0.292 |
| | Age (years) | OR = 0.93, CI = 0.72 - 1.20, p = 0.587 | | |
| | Sex assigned at birth (females) | OR =0.28, CI =0.06 - 1.14, p = 0.079 | | |
| **ID (FSIQ<=70)** | CNV | Low cell counts | | OR = 2.02, CI = 0.721 - 5.66, p = 0.181 |
| | Age (years) | OR =1.06, CI =0.88 - 1.29, p = 0.525 | | |
| | Sex assigned at birth (females) | OR =1.75, CI = 0.63 - 4.98, p = 0.282 | | |

**Supplementary Table 5.** Statistical results from linear regression analyses assessing the association between CNV group status (deletion, duplication and control) and ADI-R subdomain symptom counts.

| Outcome | Main Effect | Deletion | Duplication | Post-Hoc |
|---|---|---|---|---|
| | | | | Deletion-Duplication |
| **ADI-R SUBDOMAINS** | | | | |
| Autism - Reciprocal Social Interaction | PR-CNV Status (PR-CNV) | β =0.78, CI =0.31 - 1.24, **p =0.001** | β =1.22, CI = 0.70 - 1.73, **p =9.15x10$^{-6}$** | β =0.44, CI =-0.01 - 0.89, p =0.053 |
| | Age (years) | β =-0.01, CI =-0.08 - 0.06, p =0.782 | | |
| | Sex assigned at birth (females) | β =-0.03, CI =-0.41 - 0.35, p =0.879 | | |
| Autism (Communication and Language) | PR-CNV Status (PR-CNV) | β =0.66, CI =0.21 - 1.11, **p =0.005** | β = 1.38, CI = 0.88 - 1.88, **p =3.50x10$^{-7}$** | β =0.72, CI =0.28 - 1.16, **p =0.001** |
| | Age (years) | β =-0.01, CI =-0.08 - 0.05, p =0.681 | | |
| | Sex assigned at birth (females) | β =0.03, CI =-0.34-0.39, p =0.893 | | |
| Autism (Restricted, Repetitive, Stereotyped Behaviour) | PR-CNV Status (PR-CNV) | β =0.69, CI =0.23 - 1.15, **p =0.004** | β = 1.16, CI = 0.65 - 1.67, **p =1.73x10$^{-5}$** | β =0.47, CI =0.03 - 0.91, **p =0.038** |
| | Age (years) | β -0.05, CI =-0.12 - 0.02, p =0.160 | | |
| | Sex assigned at birth (females) | β =-0.05, CI =-0.43-0.32, p =0.777 | | |

**Supplementary Table 6.** Sensitivity analyses examining the association of blood sample collection status with psychiatric and cognitive outcomes

| Outcome | β (Successful Blood Sample) | 95% CI | p-value |
|---|---|---|---|
| **ADHD** | β = - 0.09 | CI =-0.42 – 0.23 | p=0.571 |
| **Anxiety** | β =-0.24 | CI =-0.61 – 0.13 | p=0.201 |
| **ODD** | β = 0.01 | CI =-0.40 – 0.41 | p=0.968 |
| **OCD** | β =0.01 | CI =-0.40 – 0.41 | p=0.968 |
| **Mood disorder** | β =0.02 | CI =-0.34 – 0.39 | p=0.899 |
| **Autistic Traits (SCQ>=15)** | β = - 0.11 | CI =-0.43 – 0.21 | p=0.507 |
| **DCD Traits** | β = - 0.09 | CI =-0.38 – 0.21 | p=0.559 |
| **Autism (Total ADI Score)** | β = - 0.01 | CI =-0.37 – 0.36 | p=0.959 |
| **IQ DOMAINS** | | | |
| **FSIQ** | β =4.39 | CI = - 1.12 – 9.91 | p=0.117 |
| **VIQ** | β =3.44 | CI =-2.46 – 9.34 | p=0.249 |
| **PIQ** | β =14.55 | CI =2.69 – 26.40 | **p=0.019** |

**Supplementary Table 7.** Statistical results from One-Way ANOVA and Kruskall-Wallis tests assessing genotype effects on protein levels.

| Test | Dependent variable | Main effect of genotype | Post-hoc comparisons |
|---|---|---|---|
| Kruskal-Wallis | ERK2 levels | H(2)=0.1768, P=0.9154 | |
| Kruskal-Wallis | phospho-ERK2 levels | H(2)=2.615, P=0.2705 | |
| Kruskal-Wallis | ERK1 levels | H(2)=16.50, P=0.0003 | Dunn's post-hoc: **CTR vs DEL p= 0.0245**, CTR vs DUP p=0.4791, **DEL vs DUP p=0.0003** |
| Kruskal-Wallis | phospho-ERK1 levels | H(2)=17.93, p=0.0001 | Dunn's post-hoc: **CTR vs DEL p= 0.0493**, CTR vs DUP p=0.1797, **DEL vs DUP p<0.0001** |
| Kruskal-Wallis | p70S6K levels | H(2)=13.49, P=0.0012 | Dunn's post-hoc: **CTR vs DEL p= 0.0043**, **CTR vs DUP p=0.0050**, DEL vs DUP p>0.9999 |
| One-Way ANOVA | phospho-p70S6K levels | $F_{2,53}$=3.037, p=0.0564 | |
| One-Way ANOVA | AKT levels | $F_{2,53}$=0.4541, p=0.6375 | |
| Kruskal-Wallis | phospho-AKT levels | H(2)=6.668, p=0.0357 | Dunn's post-hoc: CTR vs DEL p= 0.1657, CTR vs DUP p>0.9999, **DEL vs DUP p=0.05** |
| Kruskal-Wallis | eIF4E levels | H(2)=2.235, P=0.3271 | |
| Kruskal-Wallis | phospho-eIF4E levels | H(2)=0.2173, P=0.8970 | |
| Kruskal-Wallis | TSC1 levels | H(2)=2.856, P=0.2398 | |

**Supplementary Table 8.** Statistical results from linear regression analysis assessing the association between genotype and protein levels.

| Outcome | Main Effect | Deletion | Duplication | Post-Hoc Deletion-Duplication |
|---|---|---|---|---|
| **ERK2** | PR-CNV status (PR-CNV) | β=0.03, CI= -0.64 - 0.69, p=0.93, padj=0.97 | β=0.1, CI= -0.65 - 0.84, p= 0.79, padj=0.87 | β=-0.07, CI= -0.77 - 0.63, p= 0.84, padj=0.84 |
| | Age (years) | β=0.02, CI= -0.09 - 0.12, p=0.74 | | |
| | Sex assigned at birth (females) | β=-0.09, CI= -0.65 - 0.47, p=0.74 | | |
| **phospho-ERK2** | PR-CNV status (PR-CNV) | β=0.51, CI= -0.13 - 1.16, p=0.12, padj=0.29 | β=0.35, CI=-0.37 - 1.07, p=0.33, padj=0.51 | β=0.16, CI=-0.52 - 0.84, p=0.63, padj=0.77 |
| | Age (years) | β=0.01, CI= -0.09 - 0.11, p=0.78 | | |
| | Sex assigned at birth (females) | β=-0.19, CI= -0.74 - 0.35, p=0.48 | | |
| **ERK1** | PR-CNV status (PR-CNV) | β=-0.82, CI=-1.36 - -0.27, **p=0.004, padj=0.03** | β=0.57, CI=-0.03 - 1.18, p=0.06, padj=0.20 | β=-1.39, CI=-1.96 - -0.82, **p=1.073E-05, padj=9.7846E-05** |
| | Age (years) | β=0.02, CI= -0.06 - 0.11, p=0.55 | | |
| | Sex assigned at birth (females) | β=0.12, CI= -0.34 - 0.58, p=0.61 | | |
| **phospho-ERK1** | PR-CNV status (PR-CNV) | β=-0.78, CI=-1.33 - -0.23, p**=0.007, padj=0.04** | β=0.59, CI=-0.03 - 1.21, p=0.06, padj=0.20 | β=-1.37, CI=-1.95 - -0.79, **p=1.779E-05, padj=9.7846E-05** |
| | Age (years) | β=-0.01, CI= -0.09 - 0.08, p=0.82 | | |
| | Sex assigned at birth (females) | β=0.14, CI= -0.33 - 0.61, p=0.55 | | |
| **p70S6K** | PR-CNV status (PR-CNV) | β=-0.87, CI=-1.46 - -0.28**, p=0.004, padj=0.03** | β=-1.08, CI=-1.74 - -0.42, **p=0.002, padj=0.03** | β=0.21, CI=-0.41 - 0.82, p=0.51, padj=0.70 |
| | Age (years) | β=0.01, CI= -0.09 - 0.1, p=0.90 | | |
| | Sex assigned at birth (females) | β=-0.20, CI= -0.70 - 0.3, p=0.42 | | |
| **phospho-p70S6K** | PR-CNV status (PR-CNV) | β=-0.39, CI=-1.01 - 0.22, p=0.21, padj=0.41 | β=-0.86, CI=-1.55 - -0.16, **p=0.02,** padj=0.07 | β=0.46, CI=-0.18 - 1.11, p=0.16, padj=0.43 |
| | Age (years) | β=0.02, CI= -0.07 - 0.11, p=0.66 | | |
| | Sex assigned at birth (females) | β=-0.39, CI= -0.91 - 0.13, p=0.13 | | |
| **AKT** | PR-CNV status (PR-CNV) | β=-0.13, CI=-0.78 - 0.52, p=0.69, padj=0.84 | β=0.20, CI=-0.53 - 0.92, p=0.58, padj=0.76 | β=-0.33, CI=-1.01 - 0.35, p=0.33, padj=0.70 |
| | Age (years) | β=0.07, CI= -0.03 - 0.17, p=0.15 | | |
| | Sex assigned at birth (females) | β=0.19, CI= -0.36 - 0.73, p=0.49 | | |
| **phospho-AKT** | PR-CNV status (PR-CNV) | β=0.48, CI=-0.11 - 1.08, p=0.11, padj=0.29 | β=-0.39, CI=-1.06 - 0.27, p=0.24, padj=0.42 | β=0.88, CI=0.25 - 1.50, **p=0.007, padj=0.02** |
| | Age (years) | β=-0.09, CI= -0.18 - 0.005, p=0.06 | | |
| | Sex assigned at birth (females) | β=-0.32, CI= -0.82 - 0.18, p=0.20 | | |

| | | | | |
|---|---|---|---|---|
| **eIF4E** | PR-CNV status (PR-CNV) | β=0.38, CI=-0.27 - 1.02, p=0.24, padj=0.42 | β=0.12, CI=-0.59 - 0.84, p=0.73, padj=0.85 | β=0.25, CI=-0.42 - 0.93, p=0.46, padj=0.70 |
| | Age (years) | β=-0.06, CI= -0.16 - 0.04, p=0.23 | | |
| | Sex assigned at birth (females) | β=0.21, CI= -0.33 - 0.75, p=0.45 | | |
| **phospho-eIF4E** | PR-CNV status (PR-CNV) | β=0.23, CI=-0.43 - 0.89, p=0.49, padj=0.67 | β=-0.01, CI=-0.76 - 0.73, p=0.97, padj=0.97 | β=0.24, CI=-0.45 - 0.94, p=0.48, padj=0.70 |
| | Age (years) | β=0.003, CI= -0.10 - 0.10, p=0.96 | | |
| | Sex assigned at birth (females) | β=0.11, CI= -0.45 - 0.67, p=0.70 | | |
| **TSC1** | PR-CNV status (PR-CNV) | β=-0.40, CI=-1.03 - 0.23, p=0.21, padj=0.41 | β=-0.33, CI=-1.03 - 0.37, p=0.35, padj=0.51 | β=-0.07, CI=-0.73 - 0.59, p=0.83, padj=0.84 |
| | Age (years) | β=0.07, CI= -0.02 - 0.17, p=0.13 | | |
| | Sex assigned at birth (females) | β=-0.28, CI= -0.81 - 0.25, p=0.30 | | |

**Supplementary Table 9.** Statistical results from linear regression analysis assessing the association between symptom counts and ERK1 levels in the whole cohort (control, deletion and duplication).

| Outcome | Main Effect | Results |
|---|---|---|
| **ADHD** | ERK1 | β=0.16, CI=-0.10 - 0.42, p=0.23, padj=0.54 |
| | Age (years) | β=-0.04, CI=-0.13 - 0.05, p=0.40 |
| | Sex assigned at birth (females) | β=-0.25, CI=-0.76 - 0.27, p=0.34 |
| **Anxiety** | ERK1 | β=0.14, CI=-0.13 - 0.41, p=0.30, padj=0.55 |
| | Age (years) | β=-0.008, CI=-0.10 - 0.09, p=0.87 |
| | Sex assigned at birth (females) | β=0.26, CI=-0.27 - 0.79, p=0.33 |
| **ODD** | ERK1 | β=0.19, CI=-0.06 - 0.45, p=0.13, padj=0.54 |
| | Age (years) | β=-0.07, CI=-0.17- 0.02, p=0.10 |
| | Sex assigned at birth (females) | β=0.20, CI=-0.31- 0.71, p=0.44 |
| **OCD** | ERK1 | β=0.05, CI=-0.20 - 0.31, p=0.67, padj=0.86 |
| | Age (years) | β=0.02, CI=-0.07 - 0.11, p=0.72 |
| | Sex assigned at birth (females) | β=0.29, CI=-0.22 - 0.80, p=0.26 |
| **Mood disorder** | ERK1 | β=0.16, CI=-0.11 - 0.42, p=0.24, padj=0.54 |
| | Age (years) | β=0.02, CI=-0.07 - 0.11, p=0.68 |
| | Sex assigned at birth (females) | β=0.50, CI=-0.03 - 1.02, p=0.06 |
| **Autistic Traits (SCQ>=15)** | ERK1 | β=-0.04, CI=-0.32 - 0.24, p=0.77, padj=0.87 |
| | Age (years) | β=-0.06, CI=-0.15 - 0.04, p=0.26 |
| | Sex assigned at birth (females) | β=-0.06, CI=-0.63 - 0.51, p=0.83 |
| **DCD Traits** | ERK1 | β=-0.26, CI=-0.54 - 0.01, p=0.06, padj=0.54 |
| | Age (years) | β=-0.08, CI=-0.18 - 0.02, p=0.10 |
| | Sex assigned at birth (females) | β=-0.24, CI=-0.80 - 0.31, p=0.38 |
| **FSIQ_WASI_WISC** | ERK1 | β=-0.38, CI=-5.23 - 4.47, p=0.87, padj=0.87 |
| | Age (years) | β=0.76, CI=-0.97 - 2.49, p=0.38 |
| | Sex assigned at birth (females) | β=1.88, CI=-7.71 - 11.48, p=0.69 |
| **Autism (Total ADI score)** | ERK1 | β=0.12, CI=-0.15 - 0.40, p=0.38, padj=0.56 |
| | Age (years) | β=-0.06, CI=-0.15 - 0.04, p=0.22 |
| | Sex assigned at birth (females) | β=-0.33, CI=-0.88 - 0.22, p=0.24 |

**Supplementary Table 10.** Statistical results from linear regression analysis assessing the association between ADI-R subdomains and ERK1 levels in the whole cohort (control, deletion and duplication).

| Outcome | Main Effect | Results |
|---|---|---|
| **ADI-R SUBDOMAINS** | | |
| Autism - Reciprocal Social Interaction | ERK1 | β=0.14, CI=-0.14 - 0.42, p=0.33 |
| | Age (years) | β=-0.05, CI=-0.15 - 0.04, p=0.26 |
| | Sex assigned at birth (females) | β=-0.35, CI=-0.92 - 0.21, p=0.21 |
| Autism (Communication and Language) | ERK1 | β=0.15, CI=-0.13 - 0.43, p=0.29 |
| | Age (years) | β=-0.05, CI=-0.15 - 0.04, p=0.28 |
| | Sex assigned at birth (females) | β=-0.15, CI=-0.71 - 0.41, p=0.59 |
| Autism (Restricted, Repetitive, Stereotyped Behaviour) | ERK1 | β=0.03, CI=-0.23 - 0.30, p=0.80 |
| | Age (years) | β=-0.07, CI=-0.16 - 0.02, p=0.15 |
| | Sex assigned at birth (females) | β=-0.29, CI=-0.82 - 0.24, p=0.28 |

**Supplementary Table 11.** Statistical results from linear regression analysis assessing the association between symptom counts and phospho-ERK1 levels in the whole cohort (control, deletion and duplication).

| Outcome | Main Effect | Results |
|---|---|---|
| **ADHD** | phospho-ERK1 | β=0.22, CI=-0.03 - 0.48, p=0.08, padj=0.32 |
| | Age (years) | β=-0.03, CI=-0.12 - 0.06, p=0.47 |
| | Sex assigned at birth (females) | β=-0.25, CI=-0.76 - 0.25, p=0.31 |
| **Anxiety** | phospho-ERK1 | β=0.07, CI=-0.20 - 0.34, p=0.59, padj=0.79 |
| | Age (years) | β=-0.003, CI=-0.10 - 0.09, p=0.94 |
| | Sex assigned at birth (females) | β=0.26, CI=-0.27 - 0.79, p=0.33 |
| **ODD** | phospho-ERK1 | β=0.28, CI=0.02 - 0.53, **p=0.03**, padj=0.29 |
| | Age (years) | β=-0.07, CI=-0.16 - 0.02, p=0.13 |
| | Sex assigned at birth (females) | β=0.19, CI=-0.31 - 0.69, p=0.45 |
| **OCD** | phospho-ERK1 | β=0.04, CI=-0.21 - 0.30, p=0.72, padj=0.79 |
| | Age (years) | β=0.02, CI=-0.07 - 0.11, p=0.69 |
| | Sex assigned at birth (females) | β=0.29, CI=-0.22 - 0.80, p=0.26 |
| **Mood disorder** | phospho-ERK1 | β=0.17, CI=-0.10 - 0.43, p=0.21, padj=0.47 |
| | Age (years) | β=0.02, CI=-0.07 - 0.12, p=0.60 |
| | Sex assigned at birth (females) | β=0.49, CI=-0.03 - 1.02, p=0.06 |
| **Autistic Traits (SCQ>=15)** | phospho-ERK1 | β=0.04, CI=-0.24 - 0.31, p=0.79, padj=0.79 |
| | Age (years) | β=-0.06, CI=-0.16 - 0.04, p=0.25 |
| | Sex assigned at birth (females) | β=-0.07, CI=-0.63 - 0.50, p=0.81 |
| **DCD Traits** | phospho-ERK1 | β=-0.11, CI=-0.39 - 0.18, p=0.45, padj=0.79 |
| | Age (years) | β=-0.09, CI=-0.19 - 0.01, p=0.08 |
| | Sex assigned at birth (females) | β=-0.25, CI=-0.83 - 0.32, p=0.37 |
| **FSIQ_WASI_WISC** | phospho-ERK1 | β=0.82, CI=-4.02 - 5.65, p=0.74, padj=0.79 |
| | Age (years) | β=0.76, CI=-0.97 - 2.49, p=0.38 |
| | Sex assigned at birth (females) | β=1.80, CI=-7.79 - 11.39, p=0.71 |
| **Autism (Total ADI score)** | phospho-ERK1 | β=0.22, CI=-0.05 - 0.48, p=0.11, padj=0.32 |
| | Age (years) | β=-0.05, CI=-0.15 - 0.04, p=0.26 |
| | Sex assigned at birth (females) | β=-0.32, CI=-0.87 - 0.21, p=0.23 |

**Supplementary Table 12.** Statistical results from linear regression analysis assessing the association between ADI-R subdomains and phospho-ERK1 levels in the whole cohort (control, deletion and duplication).

| Outcome | Main Effect | Results |
|---|---|---|
| **ADI-R SUBDOMAINS** | | |
| Autism - Reciprocal Social Interaction | phospho-ERK1 | β=0.21, CI=-0.06 - 0.48, p=0.13 |
| | Age (years) | β=-0.05, CI=-0.15 - 0.05, p=0.31 |
| | Sex assigned at birth (females) | β=-0.35, CI=-0.90 - 0.21, p=0.22 |
| Autism (Communication and Language) | phospho-ERK1 | β=0.21, CI=-0.06 - 0.48, p=0.13 |
| | Age (years) | β=-0.05, CI=-0.14 - 0.05, p=0.32 |
| | Sex assigned at birth (females) | β=-0.14, CI=-0.69 - 0.41, p=0.61 |
| Autism (Restricted, Repetitive, Stereotyped Behaviour) | phospho-ERK1 | β=0.13, CI=-0.12 - 0.39, p=0.30 |
| | Age (years) | β=-0.06, CI=-0.16 - 0.03, p=0.16 |
| | Sex assigned at birth (females) | β=-0.29, CI=-0.82 - 0.23, p=0.27 |

**Supplementary Table 13.** Statistical results from linear regression analysis assessing the association between symptom counts and p70S6K levels in the combined group (control, deletion and duplication).

| Outcome | Main Effect | Results |
|---|---|---|
| **ADHD** | p70S6K | β=-0.27, CI=-0.52 - -0.01, **p=0.04,** padj=0.07 |
| | Age (years) | β=-0.02, CI=-0.11 - 0.07, p=0.62 |
| | Sex assigned at birth (females) | β=-0.26, CI=-0.76 - 0.24, p=0.30 |
| **Anxiety** | p70S6K | β=0.09, CI=-0.18 - 0.37, p=0.49, padj=0.55 |
| | Age (years) | β=-0.008, CI=-0.10 - 0.09, p=0.86 |
| | Sex assigned at birth (females) | β=0.27, CI=-0.26 - 0.81, p=0.31 |
| **ODD** | p70S6K | β=-0.16, CI=-0.42 - 0.10, p=0.24, padj=0.35 |
| | Age (years) | β=-0.06, CI=-0.16 - 0.03, p=0.18 |
| | Sex assigned at birth (females) | β=0.20, CI=-0.32 - 0.71, p=0.44 |
| **OCD** | p70S6K | β=0.09, CI=-0.16 - 0.35, p=0.47, padj=0.55 |
| | Age (years) | β=0.01, CI=-0.08 - 0.10, p=0.77 |
| | Sex assigned at birth (females) | β=0.30, CI=-0.21 - 0.80, p=0.24 |
| **Mood disorder** | p70S6K | β=-0.05, CI=-0.32 - 0.22, p=0.69, padj=0.69 |
| | Age (years) | β=0.03, CI=-0.07 - 0.12, p=0.59 |
| | Sex assigned at birth (females) | β=0.50, CI=-0.03 - 1.03, p=0.06 |
| **Autistic Traits (SCQ>=15)** | p70S6K | β=-0.43, CI=-0.72 - -0.15, **p=0.003, padj=0.03** |
| | Age (years) | β=-0.03, CI=-0.12 - 0.06, p=0.48 |
| | Sex assigned at birth (females) | β=-0.007, CI=-0.53 - 0.51, p=0.98 |
| **DCD Traits** | p70S6K | β=-0.37, CI=-0.65 - -0.09, **p=0.01, padj=0.05** |
| | Age (years) | β=-0.07, CI=-0.17 - 0.02, p=0.12 |
| | Sex assigned at birth (females) | β=-0.28, CI=-0.82 - 0.25, p=0.29 |
| **FSIQ_WASI_WISC** | p70S6K | β=5.31, CI=0.66 - 9.97**, p=0.02,** padj=0.07 |
| | Age (years) | β=0.49, CI=-1.17 - 2.16, p=0.55 |
| | Sex assigned at birth (females) | β=2.27, CI=-6.88 - 11.42, p=0.62 |
| **Autism (Total ADI score)** | p70S6K | β=-0.31, CI=-0.60 - -0.02, **p=0.04**, padj=0.07 |
| | Age (years) | β=-0.04, CI=-0.13 - 0.06, p=0.42 |
| | Sex assigned at birth (females) | β=-0.27, CI=-0.80 - 0.26, p=0.31 |

**Supplementary Table 14.** Statistical results from linear regression analysis assessing the association between symptom counts in ADI-R subdomains and p70S6K levels in the whole cohort (control, deletion and duplication).

| Outcome | Main Effect | Results |
|---|---|---|
| **ADI-R SUBDOMAINS** | | |
| Autism - Reciprocal Social Interaction | p70S6K | β=-0.33, CI=-0.63 - -0.04, **p=0.03** |
| | Age (years) | β=-0.03, CI=-0.13 - 0.06, p=0.50 |
| | Sex assigned at birth (females) | β=-0.29, CI=-0.83 - 0.25, p=0.29 |
| Autism (Communication and Language) | p70S6K | β=-0.32, CI=-0.61 - -0.02, **p=0.03** |
| | Age (years) | β=-0.03, CI=-0.13 - 0.06, p=0.52 |
| | Sex assigned at birth (females) | β=-0.08, CI=-0.63 - 0.45, p=0.75 |
| Autism (Restricted, Repetitive, Stereotyped Behaviour) | p70S6K | β=-0.16, CI=-0.45 - 0.12, p=0.25 |
| | Age (years) | β=-0.06, CI=-0.15 - 0.04, p=0.22 |
| | Sex assigned at birth (females) | β=-0.26, CI=-0.79 - 0.26, p=0.32 |

**Supplementary Table 15.** Statistical results from linear regression analysis assessing the association between symptom counts and p70S6K levels by CNV status (control versus combined deletion and duplication).

| Outcome | Main Effect | Results |
|---|---|---|
| **ADHD** | p70S6K | β=-0.05, CI=-0.30 - 0.20, p=0.70, padj=0.90 |
| | PR-CNV status (PR-CNV) | β=1.07, CI=0.54 - 1.61, **p=1.94E-04, padj=4.37E-04** |
| | Age (years) | β=0.01, CI=-0.07 - 0.09, p=0.75 |
| | Sex assigned at birth (females) | β=-0.11, CI=-0.56 - 0.33, p=0.61 |
| **Anxiety** | p70S6K | β=0.27, CI=-0.02 - 0.55, p=0.07, padj=0.59 |
| | PR-CNV status (PR-CNV) | β=0.84, CI=0.22 - 1.46, **p=0.008, padj=0.015** |
| | Age (years) | β=0.02, CI=-0.07- 0.11, p=0.69 |
| | Sex assigned at birth (females) | β=0.38, CI=-0.13- 0.90, p=0.14 |
| **ODD** | p70S6K | β=-0.08, CI=-0.37 - 0.21, p=0.58, padj=0.90 |
| | PR-CNV status (PR-CNV) | β=0.37, CI=-0.26 - 1.00, p=0.24, padj=0.27 |
| | Age (years) | β=-0.05, CI=-0.15- 0.04, p=0.29 |
| | Sex assigned at birth (females) | β=0.25, CI=-0.27- 0.77, p=0.34 |
| **OCD** | p70S6K | β=0.11, CI=-0.18 - 0.40, p=0.43, padj=0.90 |
| | PR-CNV status (PR-CNV) | β=0.10, CI=-0.52 - 0.73, p=0.74, padj=0.74 |
| | Age (years) | β=0.02, CI=-0.08- 0.11, p=0.72 |
| | Sex assigned at birth (females) | β=0.31, CI=-0.20- 0.83, p=0.23 |
| **Mood disorder** | p70S6K | β=0.03, CI=-0.27 - 0.33, p=0.85, padj=0.90 |
| | PR-CNV status (PR-CNV) | β=0.40, CI=-0.25 - 1.04, p=0.22, padj=0.27 |
| | Age (years) | β=0.04, CI=-0.06 - 0.14, p=0.43 |

| | | |
|---|---|---|
| | Sex assigned at birth (females) | β=0.55, CI=0.02 - 1.09, **p=0.04** |
| **Autistic Traits (SCQ>=15)** | p70S6K | β=-0.04, CI=-0.34 - 0.25, p=0.77, padj=0.90 |
| | PR-CNV status (PR-CNV) | β=1.35, CI=0.75 - 1.94, **p=3.52E-05, padj=1.06E-04** |
| | Age (years) | β=0.002, CI=-0.08 - 0.08, p=0.95 |
| | Sex assigned at birth (females) | β=0.08, CI=-0.36 - 0.52, p=0.72 |
| **DCD Traits** | p70S6K | β=-0.01, CI=-0.27 - 0.24, p=0.90, padj=0.90 |
| | PR-CNV status (PR-CNV) | β=1.51, CI=0.97 - 2.07, **p=1.14E-06, padj=1.02E-05** |
| | Age (years) | β=-0.02, CI=-0.10 - 0.05, p=0.57 |
| | Sex assigned at birth (females) | β=-0.09, CI=-0.52 - 0.33, p=0.66 |
| **FSIQ_WASI_WISC** | p70S6K | β=0.40, CI=-3.81 - 4.61, p=0.85, padj=0.90 |
| | PR-CNV status (PR-CNV) | β=-23.9, CI=-33.00 - -14.84, **p=2.58E-06, padj=1.16E-05** |
| | Age (years) | β=-0.29, CI=-1.67 - 1.09, p=0.68 |
| | Sex assigned at birth (females) | β=-0.92, CI=-8.45 - 6.60, p=0.81 |
| **Autism (Total ADI score)** | p70S6K | β=-0.06, CI=-0.40 - 0.28, p=0.74, padj=0.90 |
| | PR-CNV status (PR-CNV) | β=0.87, CI=0.19 - 1.55, **p=0.01, padj=0.02** |
| | Age (years) | β=-0.01, CI=-0.11 - 0.07, p=0.73 |
| | Sex assigned at birth (females) | β=-0.21, CI=-0.72 - 0.29, p=0.40 |

**Supplementary Table 16.** Statistical results from linear regression analysis assessing the association between ADI-R subdomains and p70S6K levels by CNV status (control versus combined deletion and duplication).

| Outcome | Main Effect | Results |
|---|---|---|
| **ADI-R SUBDOMAINS** | | |
| Autism - Reciprocal Social Interaction | p70S6K | β=-0.07, CI=-0.42 - 0.27, p=0.67 |
| | PR-CNV status (PR-CNV) | β=0.90, CI=0.20 - 1.59, **p=0.01** |
| | Age (years) | β=-0.01, CI=-0.10 - 0.08, p=0.83 |
| | Sex assigned at birth (females) | β=-0.23, CI=-0.75 - 0.28, p=0.37 |
| Autism (Communication and Language) | p70S6K | β=-0.10, CI=-0.45 - 0.25, p=0.58 |
| | PR-CNV status (PR-CNV) | β=0.77, CI=0.07 - 1.48, **p=0.03** |
| | Age (years) | β=-0.01, CI=-0.10 - 0.08, p=0.81 |
| | Sex assigned at birth (females) | β=-0.04, CI=-0.56 - 0.48, p=0.89 |
| Autism (Restricted, Repetitive, Stereotyped Behaviour) | p70S6K | β=0.11, CI=-0.22 - 0.44, p=0.50 |
| | PR-CNV status (PR-CNV) | β=0.96, CI=0.29 - 1.63, **p=0.005** |
| | Age (years) | β=-0.03, CI=-0.12 - 0.06, p=0.46 |
| | Sex assigned at birth (females) | β=-0.20, CI=-0.69 - 0.29, p=0.41 |

**Supplementary Table 17.** Statistical results from linear regression analysis assessing the association between symptom counts and phospho-AKT levels in the whole cohort (control, deletion and duplication).

| Outcome | Main Effect | Results |
|---|---|---|
| **ADHD** | phospho-AKT | β=-0.20, CI=-0.47 - 0.06, p=0.13, padj=0.38 |
| | Age (years) | β=0.05, CI=-0.15 - 0.04, p=0.27 |
| | Sex assigned at birth (females) | β=-0.29, CI=-0.81 - 0.22, p=0.25 |
| **Anxiety** | phospho-AKT | β=0.005, CI=-0.27 - 0.29, p=0.97, padj=0.97 |
| | Age (years) | β=-0.003, CI=-0.10 - 0.10, p=0.94 |
| | Sex assigned at birth (females) | β=0.27, CI=-0.27 - 0.81, p=0.33 |
| **ODD** | phospho-AKT | β=-0.22, CI=-0.48 - 0.05, p=0.10, padj=0.38 |
| | Age (years) | β=-0.09, CI=-0.18 - 0.005, p=0.06 |
| | Sex assigned at birth (females) | β=0.15, CI=-0.36 - 0.66, p=0.56 |
| **OCD** | phospho-AKT | β=-0.14, CI=-0.41 - 0.12, p=0.27, padj=0.54 |
| | Age (years) | β=0.005, CI=-0.09 - 0.10, p=0.91 |
| | Sex assigned at birth (females) | β=0.25, CI=-0.25 - 0.76, p=0.32 |
| **Mood disorder** | phospho-AKT | β=-0.20, CI=-0.48 - 0.07, p=0.14, padj=0.38 |
| | Age (years) | β=0.006, CI=-0.09 - 0.10, p=0.90 |
| | Sex assigned at birth (females) | β=0.45, CI=-0.08 - 0.97, p=0.09 |
| **Autistic Traits (SCQ>=15)** | phospho-AKT | β=-0.08, CI=-0.37 - 0.21, p=0.57, padj=0.79 |
| | Age (years) | β=-0.06, CI=-0.17 - 0.04, p=0.21 |
| | Sex assigned at birth (females) | β=-0.10, CI=-0.67 - 0.48, p=0.74 |
| **DCD Traits** | phospho-AKT | β=0.01, CI=-0.29 - 0.31, p=0.93, padj=0.97 |
| | Age (years) | β=-0.09, CI=-0.19 - 0.02, p=0.10 |
| | Sex assigned at birth (females) | β=-0.26, CI=-0.84 - 0.32, p=0.37 |
| **FSIQ_WASI_WISC** | phospho-AKT | β=1.34, CI=-3.68 - 6.35, p=0.59, padj=0.79 |
| | Age (years) | β=0.87, CI=-0.91 - 2.64, p=0.33 |
| | Sex assigned at birth (females) | β=2.24, CI=-7.43 - 11.9, p=0.64 |
| **Autism (Total ADI score)** | phospho-ERK1 | β=-0.08, CI=-0.37 - 0.20, p=0.55, padj=0.76 |
| | Age (years) | β=-0.06, CI=-0.16 - 0.04, p=0.21 |
| | Sex assigned at birth (females) | β=-0.34, CI=-0.91 - 0.21, p=0.22 |

**Supplementary Table 18.** Statistical results from linear regression analysis assessing the association between ADI-R subdomains and phospho-AKT levels in the whole cohort (control, deletion and duplication).

| Outcome | Main Effect | Whole group (CTR, DEL and DUP) |
|---|---|---|
| **ADI-R SUBDOMAINS** | | |
| Autism - Reciprocal Social Interaction | phospho-AKT | β=-0.10, CI=-0.39 - 0.19, p=0.49 |
| | Age (years) | β=-0.06, CI=-0.16 - 0.04, p=0.24 |
| | Sex assigned at birth (females) | β=-0.37, CI=-0.95 - 0.20, p=0.20 |
| Autism (Communication and Language) | phospho-AKT | β=-0.08, CI=-0.37 - 0.21, p=0.57 |
| | Age (years) | β=-0.06, CI=-0.16 - 0.04, p=0.27 |
| | Sex assigned at birth (females) | β=-0.16, CI=-0.73 - 0.41, p=0.58 |
| Autism (Restricted, Repetitive, Stereotyped Behaviour) | phospho-AKT | β=-0.10, CI=-0.37 - 0.17, p=0.45 |
| | Age (years) | β=-0.08, CI=-0.17 - 0.02, p=0.12 |
| | Sex assigned at birth (females) | β=-0.32, CI=-0.86 - 0.22, p=0.23 |

### References


1. Richards S, Aziz N, Bale S, Bick D, Das S, Gastier-Foster J, et al. Standards and guidelines for the interpretation of sequence variants: a joint consensus recommendation of the American College of Medical Genetics and Genomics and the Association for Molecular Pathology. Genet Med. 2015;17(5):405-24.
2. Kendall KM, Rees E, Escott-Price V, Einon M, Thomas R, Hewitt J, et al. Cognitive Performance Among Carriers of Pathogenic Copy Number Variants: Analysis of 152,000 UK Biobank Subjects. Biol Psychiatry. 2017;82(2):103-10.
3. Lynham AJ, Knott S, Underwood JFG, Hubbard L, Agha SS, Bisson JI, et al. DRAGON-Data: a platform and protocol for integrating genomic and phenotypic data across large psychiatric cohorts. BJPsych Open. 2023;9(2):e32.
4. Rammos A, Blakey R, Dennison CA, Lewis SJ, Ali N, Davies A, et al. Copy number variants and their implications for developmental and behavioural problems in cleft lip and/or palate. Hum Mol Genet. 2025;34(18):1563-74.
